# Live Imaging of muscle histolysis in *Drosophila* metamorphosis

**DOI:** 10.1101/047761

**Authors:** Yadav Kuleesha, Wee Choo Puah, Martin Wasser

**Author notes:** Corresponding Author: Martin Wasser BioImagingMW, Block 28D Dover Crescent, #31-73, Singapore 134028, Republic of Singapore.

## Abstract

**Background:** The contribution of programmed cell death (PCD) to muscle wasting disorders remains a matter of debate. *Drosophila melanogaster* metamorphosis offers the opportunity to study muscle cell death in the context of development. Using live cell imaging of the abdomen, two groups of larval muscles can be observed, doomed muscles that undergo histolysis and persistent muscles that are remodelled and survive into adulthood.

**Method:** To identify and characterize genes that control the decision between survival and cell death of muscles, we developed a method comprising *in vivo* imaging, targeted gene perturbation and time-lapse image analysis. Our approach enabled us to study the cytological and temporal aspects of abnormal cell death phenotypes.

**Results:** In a previous genetic screen for genes controlling muscle size and cell death in metamorphosis, we identified gene perturbations that induced cell death of persistent or inhibit histolysis of doomed larval muscles. RNA interference (RNAi) of the genes encoding the helicase Rm62 and the lysosomal Cathepsin-L homolog Cysteine proteinase 1 (Cp1) caused premature cell death of persistent muscle in early and mid-pupation, respectively. Silencing of the transcriptional co-repressor *Atrophin* inhibited histolysis of doomed muscles. Overexpression of dominant-negative Target of Rapamycin (TOR) delayed the histolysis of a subset of doomed and induced ablation of all persistent muscles. RNAi of *AMPKα,* which encodes a subunit of the AMPK protein complex that senses AMP and promotes ATP formation, led to loss of attachment and a spherical morphology. None of the perturbations affected the survival of newly formed adult muscles, suggesting that the method is useful to find genes that are crucial for the survival of metabolically challenged muscles, like those undergoing atrophy. The ablation of persistent muscles did not affect eclosion of adult flies.

**Conclusions:** Live imaging is a versatile approach to uncover gene functions that are required for the survival of muscle undergoing remodelling, yet are dispensable for other adult muscles. Our
approach promises to identify molecular mechanisms that can explain the resilience of muscles to PCD.

## Background

Skeletal muscles are essential for mobility and metabolism. Therefore, the preservation of muscles mass and strength plays an important role in improving the quality of life in sickness and old age. The two most common types of muscle wasting are sarcopenia, the ageing related loss of skeletal muscles, and cachexia, a metabolic syndrome associated with diseases such as cancer, heart failure and HIV [1]. Muscle degeneration can also result from heritable muscular dystrophies, such as Duchenne muscular dystrophy [2] or centronuclear myopathies [3].

Muscle wasting can result from two cellular processes; the reversible reduction of muscle fiber size by atrophy or the irreversible elimination of muscles by programmed cell death (PCD). The changes of muscle size are controlled by conserved signaling pathways that regulate the rates of protein synthesis and degradation. Protein synthesis and cell growth are activated by a signaling cascade consisting of insulin-like growth factor-1 (IGF-1), the kinase Akt1 and the mammalian target of rapamycin (mTOR) [4]. Protein breakdown and atrophy are activated by a pathway comprising Myostatin, Smad2 and the FoxO transcription factors [5]. Proteins are degraded by two processes, the autophagy lysosomal pathway [6] and the ubiquitin proteasome system [7].

Less understood is the contribution of PCD to muscle wasting. Satellite cells are myogenic stem cells that fuse with muscles to repair injuries and drive hypertrophy. In old age, satellite cells are more prone to apoptosis, thus promoting sarcopenia by affecting the repair of damaged muscles [8]. The relationship between cell death in postmitotic, multinucleated muscle fibers and muscle wasting is less clear. Skeletal muscles are considered to be more resistant to PCD than proliferating and mono-nucleated cells. In cultured C2C12 cells treated with the apoptosis inducers H_2_O_2_ or staurosporine, apoptosis was significantly reduced to 10% in myotubes compared to 50% in myoblasts, although several pro-apoptotic proteins like caspases, EndoG and AIF were induced [9]. The protective effect in differentiated muscles was proposed to be mediated by yet undefined anti-apoptosis mechanisms. Muscle fibers exhibit morphological changes during PCD that are different from those observed in mono-nucleated cells. The structural analysis of rat skeletal muscles during post-denervation atrophy revealed cell shrinkage and chromatin condensation. However, the occurrence of DNA fragmentation, a hallmark apoptotic phenotype, was negligible [10]. Furthermore, apoptosis associated with atrophy in rat muscles was found outside of but not inside muscle fibers [11].

Most knowledge about PCD in mammalian muscles is derived from experimental interventions like denervation, immobilization or exposure to toxins. As such, progress is hampered by the lack of models to study naturally occurring muscle cell death. Nonmammalian models can fill this knowledge gap since degeneration of skeletal muscles can be studied without experimental manipulations during amphibian and insect development. In *Xenopus laevis* metamorphosis, thyroid hormone induces apoptosis of tail muscles in the tadpole[12], which is associated with the formation of muscle apoptotic bodies (sarcolytes) and chromatin fragmentation. During metamorphosis of *Lepidoptera* (moths and butterflies), most muscles of caterpillars are destroyed, while a few persistent muscles survive into adulthood [13]. Major insights in moths were obtained from research on the giant intersegmental muscles (ISM) of the abdomen that undergo hormonally induced atrophy prior to eclosion and die after eclosion [14]. ISM PCD, which is triggered by changes in juvenile hormone 20-hydroxyecdysone, does not display a variety of morphological features associated with classical apoptosis, such as chromatin condensation or phagocytosis [16, 17].

Metamorphosis of the fruit *Drosophila melanogaster* is another insect model to study apoptosis of muscles in the context of animal development. Compared to the moth models, it offers an arsenal of genetic tools such as targeted reporter gene expression and gene perturbation. Since the pupal cuticle is transparent, muscle development can be followed by live cell imaging. Skeletal muscles arise in embryogenesis through the fusion of founder cells with fusion competent myoblasts [18]. During the larval stages lasting 4 to 5 days, body wall muscles grow up to 50-fold in size while their number stays constant [19]. During the subsequent 5 days of metamorphosis, which transforms larvae into adult flies, larval muscles follow two major developmental pathways. Most muscles undergo cell death and break down into sarcolytes. A second population of persistent muscles is resistant to histolysis and survives to adulthood. The alternative fates can be observed simultaneously in the pupal abdomen by live imaging of muscles marked with fluorescent proteins [20]. Dorsal external oblique muscles (DEOMs) disintegrate coincident with head eversion (HE) at the prepupal to pupal transition (PPT), which takes place around 12 hours after puparium formation (Figure 1A). More basally located dorsal internal oblique muscles (DIOMs) are resistant to pupal cell death and are remodeled into temporary adult muscles, that will later degenerate within 24 hours of eclosion [21]. Remodeling of DIOMs involves atrophy in early and growth in late metamorphosis. The molecular mechanisms regulating PCD of larval muscles remain poorly understood. Ecdysone receptor signaling activates cell death of muscles and other larval tissues. Cell death in midgut and other tissues is promoted by the autophagy lysosomal pathway [22]. However, autophagy does not appear to contribute to cell death of muscles. Instead, histolysis of muscles appears to involve apoptosis [23]. Even less is known about the genes that protect persistent muscles from hormone-induced histolysis.

**Figure 1.**
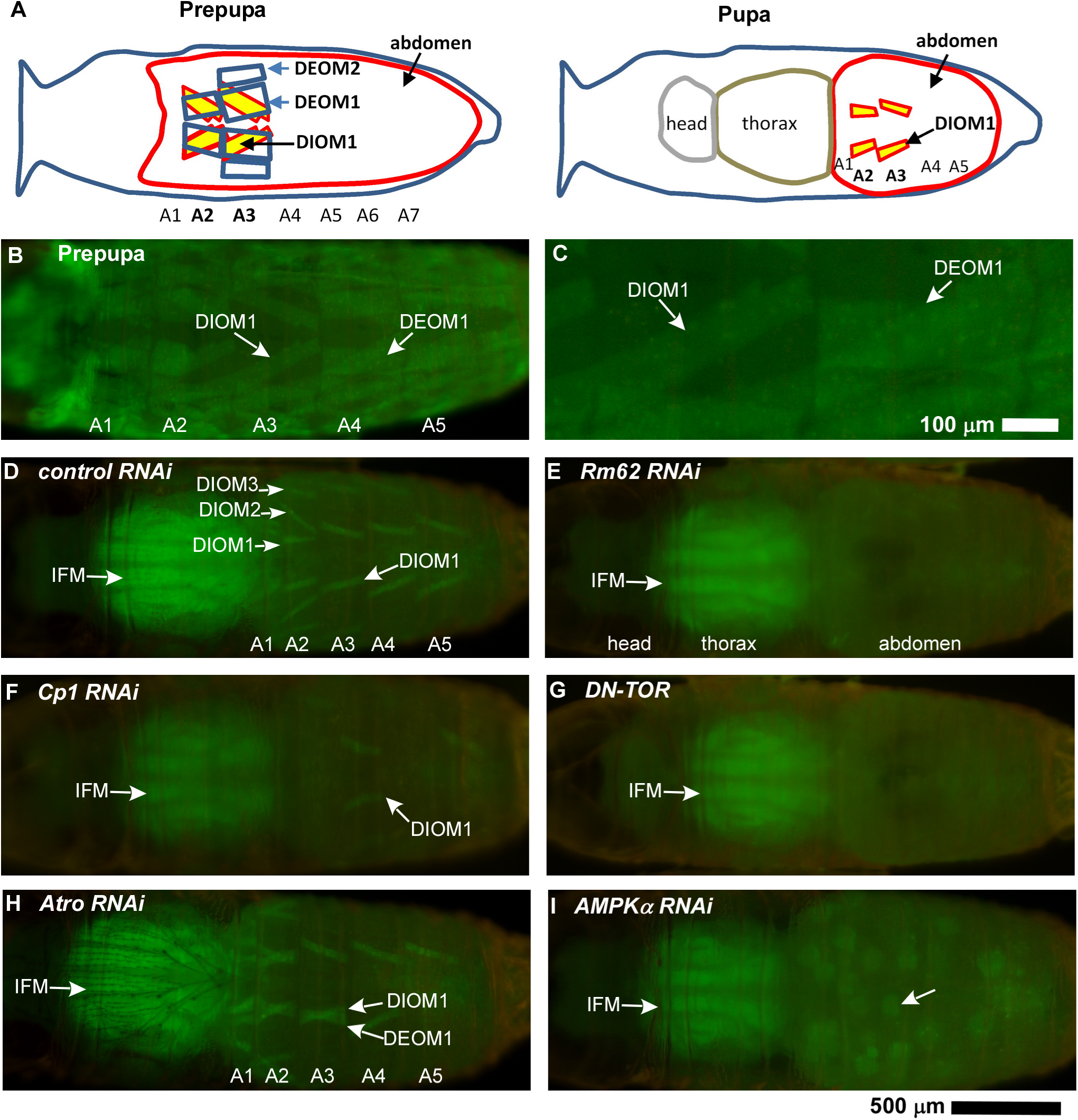
Detection of muscle cell death phenotypes by macro zoom microscopy. (A)Anatomical drawings of the dorsal sides of a fly during prepupal and pupal stage illustrate the fates of two population of larval body wall muscles. While the apically located DEOMs (blue) are destroyed during and right after head eversion, the more basally located DIOMs (yellow) survive throughout pupation into adulthood. A1-A7 indicate the positions of abdominal segments. Panels (B-I) show dorsal views (anterior on the left) of a prepupa (B,C) and late pupae (D-I) expressing the *MHC-tau-GFP* reporter to label muscles that were acquired using a macro zoom microscope The first five abdominal segments (A1-A5) are indicated. (B) In prepupae, live doomed muscles like a DEOM1 in A4 and persistent muscles like a DIOM1 in A3 can be viewed through transparent cuticles. (C) Close-up of (B). (D) In late pupation, persistent larval muscles that escaped histolysis and were remodeled are seen in A1-A5. The thorax contains the prominent indirect flight muscles (IFM). (E) RNAi of *Rm62*, (F) RNAi of *Cp1* and (G) overexpression of dominant negative TOR (*TOR*^TED^) induced cell death of persistent muscles. (H) *Atrophia (Atro)* RNAi prevented histolysis of doomed DEOMs in the 2^nd^ and, occasionally, 3^rd^ abdominal segment. (G) *AMPKα* RNAi induced degeneration of DIOMs into muscle spheroids (arrow). None of the gene perturbation affected the development of newly formed adult muscles like the IFMs. The 500 μm bar corresponds to all panels except (C).

In previous studies, we demonstrated that *in vivo* microscopy could image abdominal muscle development throughout the 5 days of *Drosophila* metamorphosis [20, 24]. Here we extend our methodology to identify and characterize genes that are involved in the PCD of doomed muscles and the survival of remodeled muscles. We used a macro zoom microscope to screen for candidate gene perturbations that either prevent or induce cell death, while multi-location laser scanning confocal microscopy (LSCM) helped us to elucidate their phenotypic consequences in more detail. Loss of function of *TOR*, the RNA helicase *Rm62*, the lysosomal protease cathepsin-L homolog *Cp1* and the master regulator of energy metabolism *AMPKα* caused histolysis of persistent larval muscles. However, none of these gene perturbations affected the survival of newly formed adult muscles. In contrast, reducing the expression of *Atrophin* inhibited histolysis of a subset of doomed larval muscles. Since this approach was able to identify genes that protect Drosophila persistent muscles from degeneration, it is conceivable that some conserved genes may also promote resilience of mammalian skeletal muscles to atrophy.

## Methods

### Drosophila Stocks

We used the UAS-GAL4 system [25] for targeted expression of fluorescent reporter genes, small hairpin (sh) RNAs and effector proteins in muscles. *P*{*Mef2-GAL4*} on the 3^rd^ chromosome (B-27390, Bloomington Drosophila stock center) served as a muscle-specific driver [26] and was recombined with *P{UAS-His2av-mKO}3* (B-53731) to mark myonuclei with Histone H2Av-mKO (monomeric kusabira orange), henceforth referred to as histone-mKO. *Mef2* is the *Drosophila* homolog of the vertebrate *myocyte-specific enhancer factor 2*, which is expressed in mesoderm, visceral and somatic muscles [27]. *P*{*MHC-tau-GFP*}1 was used to label the cytoplasm of muscle fibers. The GFP-reporter linked to the promoter of the *myosin heavy-chain (MHC)* gene is expressed in somatic musculature [28]. All UAS-shRNA strains were derived from Transgenic RNAi Project (TRiP) collection [29]. The UAS-shRNA and UAS-effector transgenic lines affecting cell survival and death of larval abdominal muscles studied in this report are listed in Table 1. A complete list of the UAS-lines used for gene perturbations in out pilot screen was previously described [30] and is provided as a supplement (Additional file 1: Table S1). In addition, we knocked down *TOR* by RNAi using the stock B-35578 (TRiP# GL00156). The *P*{*UAS-Cp1-mKO2*} reporter line [31] was used to evaluate the target specificity of Cp1-sRNA constructs. To visualize cell death in live muscle and non-muscle cells, we used the enhancer trap line P{GawB}how[24B] on the 3^rd^ chromosome (B-1767) that was recombined with *P*{*UAS-His2av-mKO*} and crossed with the mitochondrial marker *P*{*UAS-Mito-HA-GFP*}3 (B-8443)[32] to visualize the cytoplasm.

**Table 1.**
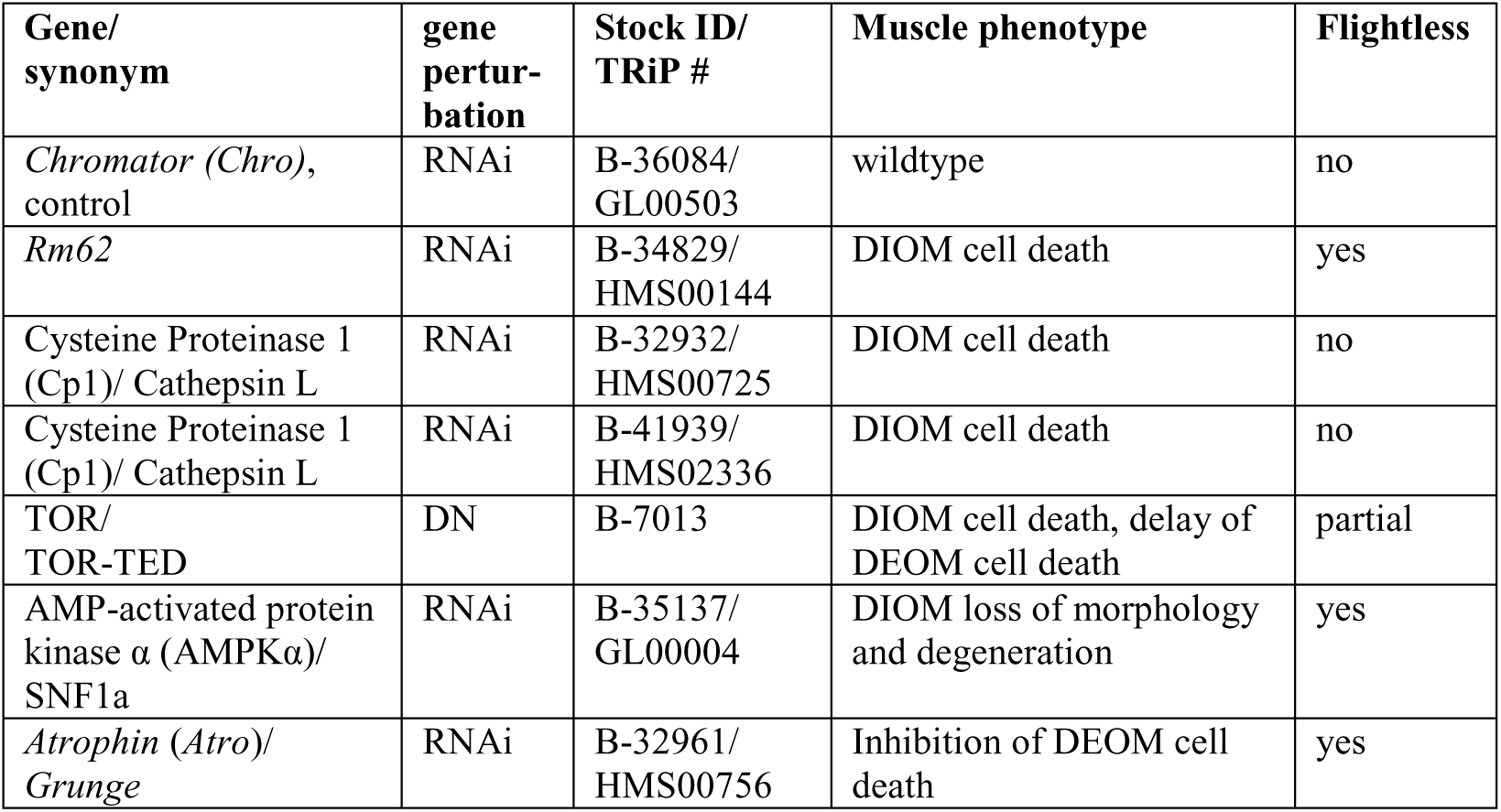
List of gene perturbations that affect survival of larval abdominal muscles in metamorphosis. Initial phenotypic assessment of a minimum of 20 pupae per genotype was carried out using a macro zoom microscope.

In our gene perturbation experiments, we crossed females of the reporter line *MHC-tau-GFP/FM7-GFP; Mef2-GAL4, UAS-histone-mKO/TM6B Tb* with males of the *UAS-GeneX-cDNA* or *UAS-GeneX-shRNA* lines. From the progeny, we selected non-*Tubby* prepupae expressing both fluorophores (e.g. MHC-tau-GFP/+; Mef2-GAL4, UAS-histone-mKO/UAS-GeneX-shRNA) for inspection of muscle phenotypes. For convenience, we will refer to these animals as Muscle-GO-GP (GO=Green+ Orange live reporter, GP=Gene perturbation). The samples were examined using an Olympus MVX10 fluorescence macro zoom microscope (Olympus, Japan). The *UAS-Chro-shRNA* construct (TRiP#GL00503, B-36084) when crossed with Muscle-GO-GP displayed no abnormalities of muscles development, eclosion and ability to fly and was therefore used as control throughout this study.

### Screening for muscle phenotypes using macro zoom microscopy

Usually, 20 Muscle-GO-GP prepupae were arranged dorsal side up in 4 groups of 5 samples on plastic Petri dishes. We recorded images at daily intervals for up to 6 days or until the adult flies eclosed using an Olympus MVX10 macro zoom microscope equipped with a DP73 digital CCD camera and cellSens acquisition software. Macro zoom microscopes are equipped with zoom objective lenses of high numerical aperture (N.A.) to resolve fine details in large samples at long working distances. We used the MVPLAPO 1× objective lens with an N.A. of 0.25. The fields of view were recorded twice with filters for green and orange fluorescence, at a zoom factor of 1.25, all of which resulted in digital color images (TIF or PNG format) of 2400×1800 pixels and a pixel size of 2.41 microns/pixel. Images were stored on a shared network drive to facilitate the visual inspection of phenotypes by multiple observers. To assess the effects of gene perturbations on muscle function we monitored eclosion and the ability to fly in adult flies.

### Time-lapse confocal microscopy of metamorphosis

The protocol for sample preparation and time-lapse imaging of *Drosophila* pupae was previously described [33]. Live samples were collected at the white pupal stage, rinsed with water to remove the fly food from their surface and inspected under a macro zoom fluorescence microscope to confirm expression of both reporter genes. Up to 30 prepupae were positioned dorsal side down on an uncoated 32 mm diameter glass bottom dish (MatTek, Ashland, Massachusetts). The live samples were mounted in CyGEL (Biostatus Ltd, Leicester, UK) to restrict their movement during imaging. Head eversion leads to compression and a posterior shift of the abdomen. As our goal was to view pupal abdominal segments 1-5 during live imaging, prepupae had to be placed in such a way that the anterior border of their 3^rd^ abdominal segment was adjacent to one side of the field of view. A wet tissue was kept around the specimens to maintain humidity levels during imaging. We used the Zeiss LSM 5 Live (Carl Zeiss, Jena, Germany) inverted line scanning confocal microscope equipped with a motorized XY scanning stage to perform multi-location timelapse imaging. 3D time-lapse image acquisition was carried out for 5 days at 30 minute intervals (240 time points per sample) using a 10×/0.3 EC-Plan-Neofluar M27 air objective, at a scan zoom of 0.5. The two colour channels were recorded sequentially; channel1 with an excitation laser of 488 nm, band path (BP) filter 500-525; channel 2 with 532 nm laser line, BP 560-675. Image stacks containing 35-40 optical slices were collected at 13.2 μm intervals. Each optical slice had a frame size of 1024 × 1024 pixels with a pixel size of 1.25 μm. The manufacturer’s Zen 2008 software was used for image acquisition, with the built-in multi-time series (MTS) macro controlling repetitive scanning of multiple locations. The MTS saved one LSM image file per time point and location. The temperature of the microscope room was set to 22°C for most experiments. LSCM recordings of samples expressing *Atro* and *AMPKα* shRNA were performed at 25°C. We used a data logger device to record temperature and humidity during imaging. The speed of metamorphosis depends on temperature Samples imaged by LSCM at 22°C eclosed on average 107 hours after entering head eversion, while this period was reduced to around 75 hours at 25°C.

### Image Analysis Workflow

We previously introduced a pipeline for the visualization and quantification of *in vivo* microscopy data [33]. Most steps were carried out using custom software tools, unless otherwise indicated. Using the TLM-Converter custom software [34], we concatenated the image stacks stored in 8-bit LSM format to create one 3D time-lapse ICS file per sample with sizes ranging from 17 to 19 Gigabytes (GB). 3D stacks in ICS format were converted to maximum intensity projections (MIPs) to generate 2D time-lapse images which were saved as multi-page TIFF files. Uncompressed TIFF files of 240 time points had sizes of 737 MB. TIFF files could be compressed over 20 fold using JPEG compression without noticeable degradation in image quality.

Phenotypic analysis of time-lapse image data was performed using the TLM-Explorer tool [24]. Besides experimental parameters such as genotype, time interval and resolution, the user has to define the onset of head eversion (HE) as the temporal reference point for comparing different datasets. HE marks the transition from the larval to the tripartite adult body plan comprising head, thorax and abdomen. The beginning of HE is recognized by rapid movements of abdominal muscles that lead to misalignments between optical slices during 3D imaging and blurring of image projections. Developmental time points are indicated in hours and minutes (h:m) after head eversion (aHE), where negative values refer to the prepupal and positive to the pupal stage. In addition, user can draw ROIs to measure cell size and visualize changes of nuclear morphology inside cell boundaries.

The workflow for morphological quantification was implemented as a custom tool in the C++. NET framework and was named QuaMMM (Quantitative Microscopy of Muscles in Metamorphosis). We used the following libraries: FreeImage [35] handled the processing of multi-page TIFF files and libics the import of ICS files [36]. To prepare videos, we exported annotated frames as JPEG files and built animations using the VirtualDub video processing software. Videos were exported as AVI files and converted to MP4 using the Freemake Video converter software. To create the figures in our manuscript, we used Photoshop CS3 (Adobe) and ACDSee Pro 5 (ACD Systems Int. Inc.).

### Statistical Analysis

Statistical data analysis was performed using Excel (Microsoft) and Minitab 16 (Minitab Inc.). Minitab was used to compute the confidence intervals of proportions and produce box-and-whisker plots (boxplots). Excel was applied to calculate proportions and plot bar charts.

## Results

### Live imaging based screening for genes regulating survival of larval muscles in metamorphosis

Fluorescent microscopy reveals the contrasting fates of dorsal abdominal muscles in prepupae. Dorsal external oblique muscles (DEOMs) are eliminated by programmed cell death, while the dorsal internal oblique muscles (DIOMs) resist histolysis and persist into adulthood (Figure 1A-C). According to nomenclature in the literature [21], the most dorsal muscles in the first five abdominal segments A1 to A5 are referred to as DEOM1 and DIOM1, more lateral ones as DEOM2/3 or DIOM2/3. To identify genes that promote cell death of doomed or survival of persistent larval muscles during metamorphosis, we developed a screen that is based on live imaging, targeted gene perturbation and time-lapse image analysis. Females of a master stock containing three transgenes, the muscle specific driver *Mef2-GAL4, UAS-histone-mKO* to label nuclei and *MHC-tau-GFP* to visualize the cytoplasm of muscles, were crossed to males carrying UAS effector constructs to drive the expression of transgenic proteins and small hairpin (sh) RNAs for RNA interference (RNAi). The phenotypic effects of muscle specific gene perturbation were assessed by *in vivo* microscopy in two steps as previously described [31]. First, to screen for interesting phenotypes, we monitored live muscle development in a minimum of 20 specimens per genotype at daily intervals using a fluorescence macro zoom microscope. To evaluate effects on muscle function, we scored eclosion rates and the ability to fly. Second, gene perturbations resulting in interesting phenotypes were further examined by 3D time-lapse LSCM from the prepupal to pharate adult stage for 4-5 days at 30 minute intervals. Timelapse imaging was performed in multiple locations (up to 30 samples) using a x-y scanning stage. Using custom software, we created time-lapse maximum intensity projections (MIPs) of 3D image stacks and performed time-series image analysis of muscles development during metamorphosis. We had earlier performed a pilot screen with 119 publicly available UAS fly stocks, involving 98 unique genes, which were targeted by 100 RNAi and 19 protein-overexpression constructs [30] (Additional file 1: Table S1). Five of the gene perturbations specifically interfered with cell death or survival of larval muscles without affecting the development of newly formed adult muscles (Table 1). Macro zoom microscopy revealed that RNAi of the RNA helicase *Rm62* (Figure 1E) and *Cysteine proteinase 1 (Cp1)* (Figure 1F), and overexpression of dominant negative *TOR* induced the removal of DIOMs (Figure 1G).*Atrophin* RNAi prevented histolysis of a subset of DEOMs (Figure 1H). RNAi of AMP-activated protein kinase α subunit (*AMPKα*) caused the loss of tubular morphology and degeneration of DIOMs (Figure 1I). About half of the RNAi constructs displayed wildtype morphology and function of muscles, which were indistinguishable from flies without RNAi overexpression. One of these constructs, UAS-Chro-shRNA, was chosen as a control.

### Live imaging of muscle histolysis by confocal microscopy

The dynamics of muscle histolysis can be viewed in detail at the subcellular level using LSCM (Figure 2). Prior to head eversion (HE), DEOM1s lose their rectangular shape and display a curvy contour (Figure 2A, 2B), while nuclear fluorescence increases due to chromatin condensation (Figure 2A’, 2B’). More laterally located DEOM2s undergo histolysis after HE, usually between +5h to +10h aHE (Figure 2C, 2D). Once the vigorous muscle contractions during HE have stabilized, DEOM1s are already shattered and continue to disintegrate into sarcolytes that retain tau-GFP and, occasionally contain condensed myonuclei labelled with histone-mKO (Figure 2C-H, arrow heads). Muscle histolysis does not unfold like a typical apoptosis. Nuclei condense without subsequent fragmentation. In fact, bright condensed nuclei can be observed until late pupation (Figure 2G), suggesting negligible, if any, nuclear fragmentation occurs. The difference between PCD in muscles and mono-nucleated cells (MNCs) could be visualized using the muscle-specific Gal4-driver 24B-Gal4 which is expressed in muscles and more apically located MNCs, presumably hypodermis (Additional file 2: Figure S1). After +6h aHE, nuclei of MNCs condensed and showed fragmentation (Figure 2I). We determined an average duration between nuclear condensation and fragmentation of 45.4 ± 23.2 minutes (2 pupae; 14 cells; 95% CI of mean 32.0 to 58.7). In contrast, we did not observe any fragmentation of condensed myonuclei in dying muscles of the same pupae (Figure 2J). The absence of nuclear fragmentation is consistent with studies on Manduca and denervated rat muscles, all of which support the idea that PCD in muscles is different from classical apoptosis [10, 17]. Fluorescent proteins in sarcolytes showed remarkable stability, persisting up 3-4 days following HE, indicating that membranes of sarcolytes remained intact to prevent leakage and that proteolysis was subdued. Muscle fragmentation, instead of rapid self-destruction, may help maintain the role of muscles as a reservoir of amino acids and other metabolites that can be slowly released during metamorphosis.

**Figure 2.**
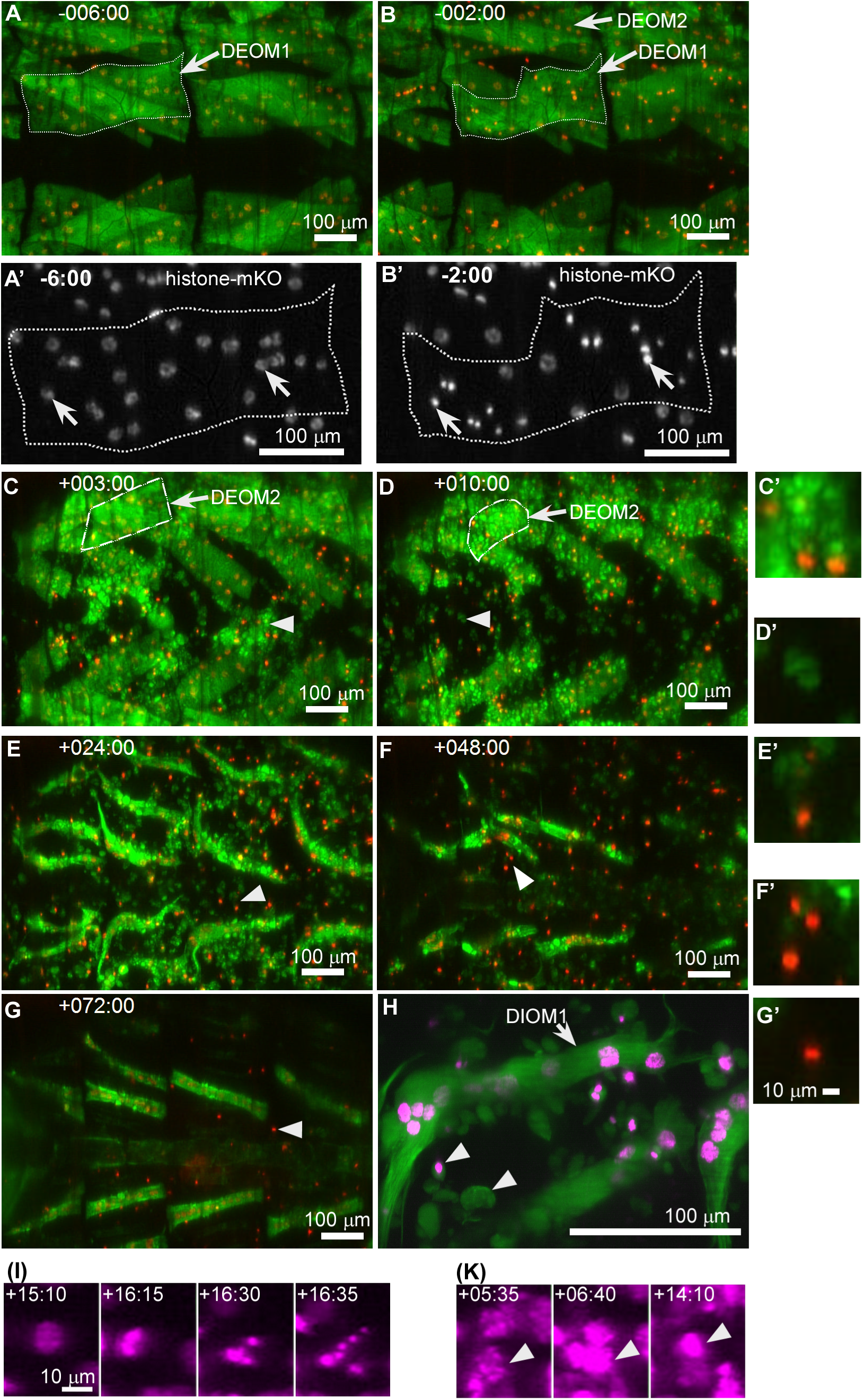
Cell death in wildtype doomed muscles observed by confocal microscopy. All images show dorsal views (anterior left) of prepupal (A, B) and pupal (C-H) abdominal regions. Muscles were labeled with Mhc-tau-GFP (green) and myonuclei with UAS-histonemKO (red, except for A’, B’ in white and H, I, K in magenta) driven by Mef2-Gal4. (A-G) correspond to the same specimen. (A, B) Disintegration of DEOM1s is initiated prior to HE and accompanied by condensation of myonuclei (A’, B’). Muscles outlined in (A,B) were magnified two-fold in panels (A’/B’). (C-H) Doomed muscles break apart into sarcolytes (arrow heads) which contain tau-GFP and, occasionally, condensed nuclei. (C, D) More laterally positioned DEOM2s undergo histolysis after completion of HE. (H) High magnification image of a DIOM1 located in the left hemi-segment of A2 on the second day of pupation surrounded by fluorescently labeled sarcolytes that persist for up to 3 days. (I, K) UAS-histone-mKO labeled cells driven by 24B-Gal4. (I) PCD of mono-nucleated, non-muscle cells (see Additional file 2: Figure S1) is associated with nuclear fragmentation following nuclear condensation. (K) In contrast, muscle fibers like the DEOM2 show condensation but not fragmentation of nuclei.

### DIOMs are not required for eclosion

As previously reported, DIOMs are eliminated in adult flies on the first day after eclosion [21]. This temporary nature suggested that these remodeled abdominal muscles may only be required during eclosion and become obsolete in adult flies. Unexpectedly, partial or total loss of DIOMs resulting from *Cp1* or *AMPKα* RNAi, resulted in near-wildtype eclosion rates (Figure 3). Therefore, the function of DIOMs in late metamorphosis and the purpose of their elaborate remodeling remain to be elucidated.

**Figure 3.**
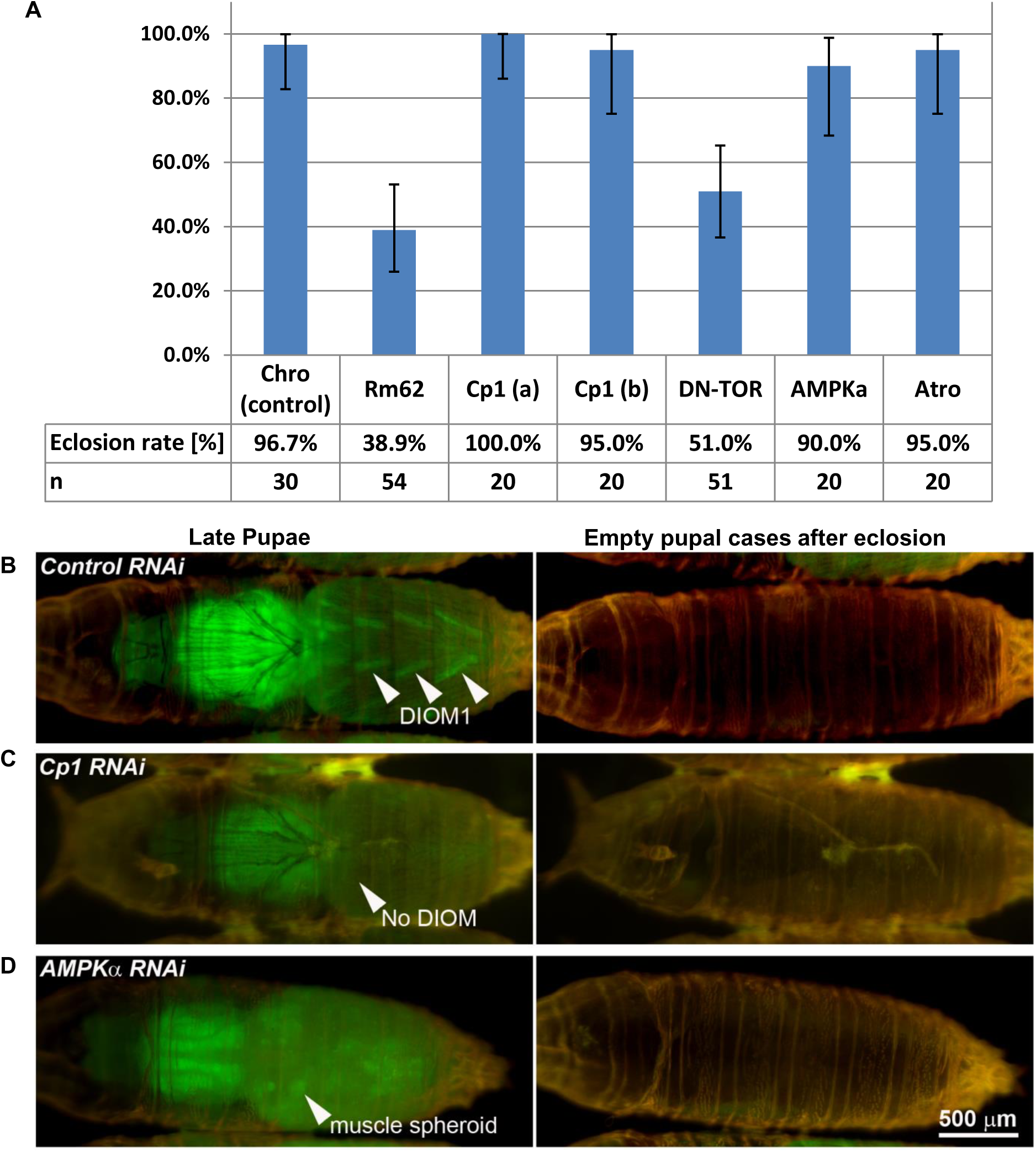
DIOMs are not required for eclosion. (A) Eclosion rates were scored for the indicated number of samples n per genotype. Error bars indicate the 95% confidence intervals of the proportions. (B-D) Images of late pupae (right column) and their cases after eclosion (left column) expressing the indicated RNAi constructs in muscles using the Mef2-Gal4 driver. Muscles were labelled with Mhc-tau-GFP (green). Shells were visualized by auto-fluorescence. (A) Arrow heads of a control pupa indicate DIOM1s. (B) Although overexpression of *Cp1-shRNA* induced elimination of all persistent muscles, the adult fly managed to eclose. An arrow head points at abdominal segment A2. (C) Similarly, the conversion of DIOMs to muscle spheroids (arrow head) and their subsequent elimination in response to AMPKα RNAi did affect the eclosion of the adult fly.

### Validation of RNAi experiments

All UAS-shRNA constructs used were generated by the Harvard Transgenic RNAi Project (TRiP), which applied bioinformatics methods in the selection of target sequences to avoid off-target effects [29]. The following lines of evidence support our notion that the shRNA constructs that caused cell death related phenotypes silenced the intended targets. Cell death by Cp1 silencing could be induced with two different shRNA constructs targeting different regions of the *Cp1* mRNA (Table 1). Using a previously described fluorescent reporter gene expressing Cp1-mKO2 [31], we were able to show that both Cp1-shRNA were able to abolish muscular fluorescence in prepupae and pupae (Additional file 2: Figure S2).

A previous study reported that the same *AMPKα* shRNA construct used in this study reduced *AMPKα* mRNA expression in embryos by 95% when expressed using the maternal MTD-Gal4 driver [38]. A previous genome-wide muscle-specific RNAi screen showed that *Rm62* and *Atro* shRNAs constructs from the Vienna Drosphila RNAi Center (VDRC), which were expressed using the same Mef2-Gal4 driver used in this study, induced flightlessness, an indicator of adult muscle defects [26]. The *Rm62* and *Atro* RNAi constructs of the VDRC and TRiP collections target different regions of the corresponding mRNAs. In addition, consistent with our results, other studies have shown that *Atro* overexpression contributed to cell death in the nervous system [39] and was involved in embryonic myogenesis [40].

### TOR and Rm62 protect persistent muscles from histolysis

Ecdysone receptor signaling activates cell death of DEOMs [23] and most other doomed larval tissues. However, little is known about the factors that protect persistent muscles from programmed cell death during metamorphosis. Our live imaging approach helped to identify three gene perturbations, *Rm62* RNAi, *Cp1* RNAi and *TOR*^TED^ overexpression, that caused premature cell death of DIOMs, yet did not affect survival of newly formed adult muscles such as IFMs, heart or abdominal muscles (Figure 4). To quantify premature cell death in persistent muscles, we scored rate and time of death (TOD) of the six DIOM1s in abdominal segments A2, A3 and A4 imaged by LSCM at 22 °C (Figure 5). Cell death rate was defined as the proportion of DIOM1s that disintegrated during pupation, while TOD records the period during pupation, when cell morphology becomes distinctly different from control muscles, e.g. change from straight to curvy edges and loss of tubular morphology. (Addional file 4: Video 1). Despite their common terminal phenotypes, the gene perturbations differed in TOD and morphological changes during histolysis. In control animals, DEOM1s began to disintegrate prior to prepupal to pupal transition (PPT) and gave rise to fluorescently labeled sarcolytes, (Figure 4A; Additional Video 1). The overexpression of dominant negative TOR (DN-TOR), also referred to as TOR™ (toxic effector domain) comprising amino acids 1228-1947 of the 2470 residue long TOR kinase [33], caused complete elimination of DIOMs in the period from +5 to +20 h aHE (Figure 4B). Unlike in controls, the DEOM1s persisted through PPT. At +4h, 89.6% of 48 DEOM1s (95% CI: 77.3%, 96.5%) in A2 to A4 of 8 *TOR*^TED^ pupae were still intact. At +5 h, both DEOMs and DIOMs started to change morphology and condense. Subsequently, both types of muscles disintegrated to give rise to sarcolytes that were noticeably larger than in controls (compare muscle fragments in Additional file 1: Video 1). TOR^TED^ has been proposed to act as a dominant-negative inhibitor of endogenous TOR-kinase that induces autophagy [41] However, silencing of *TOR* by RNAi, which caused highly penetrant and significant atrophy of DIOMs, led to negligible cell destruction [31]. Cell death rate resulting from TOR-RNAi was 12.8% with a mean TOD of +25.6h compared to 100% DIOM histolysis for TOR-TED with a mean TOD of +6.2h (Figure 5). The phenotypic differences suggest that TOR^TED^ does not behave like a *TOR* loss-of-function allele in the context of DIOM development.

**Figure 4.**
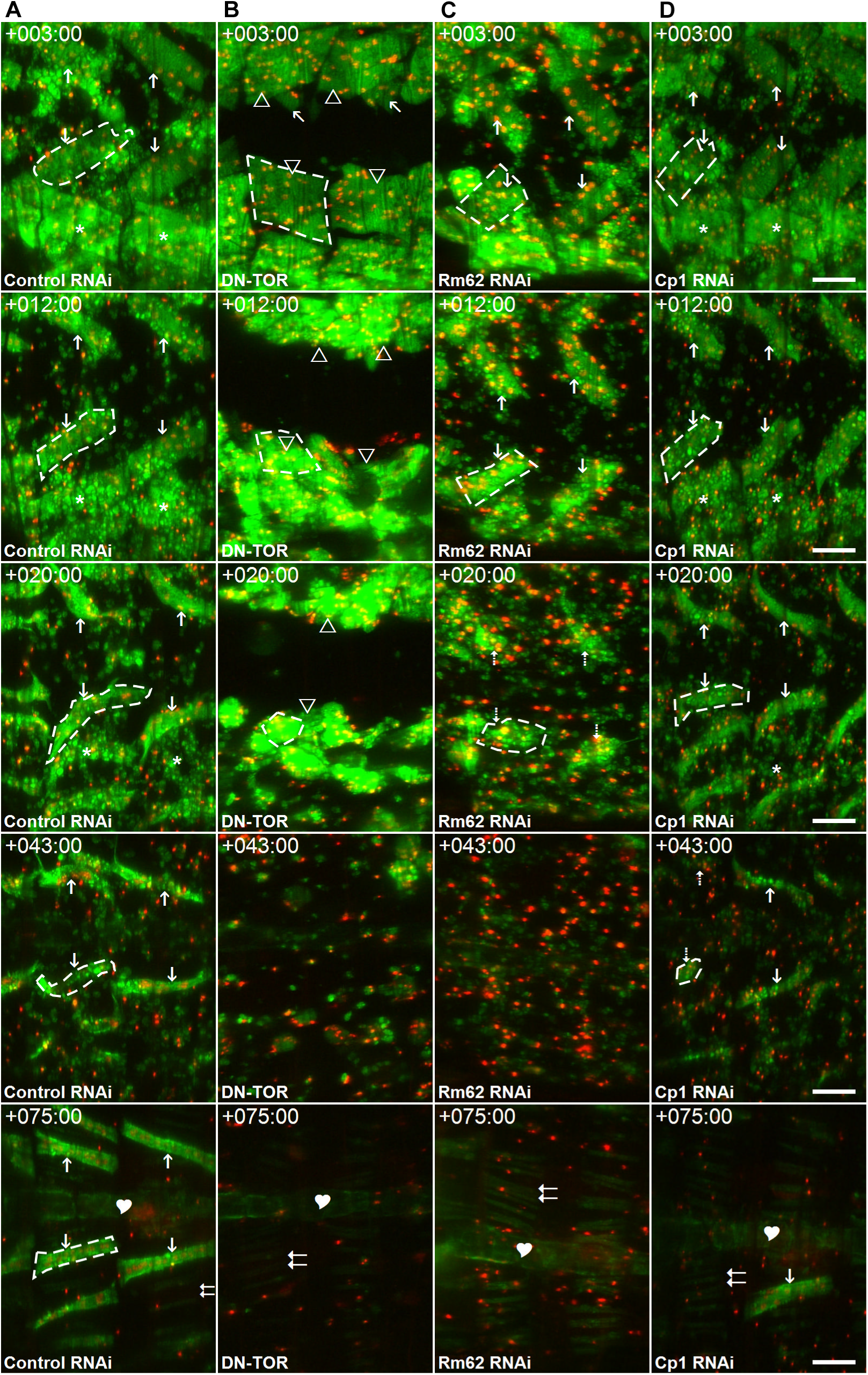
Identification of genes required for the survival of persistent muscles. Dorsal views of abdominal segments A2 and A3 of four genotypes are shown in five time points after head eversion in the anterior-posterior orientation from left to right. Muscles were labelled with tau-GFP (green) and histone-mKO (red). For more details, see Additional Video 1.(A) In a pupa expressing a control shRNA, persistent muscles (DIOM1) near the midline (vertical arrows) survived to adulthood, while laterally located obsolete DEOM2s (*) were fragmented into fluorescent sarcolytes that persisted into late pupation. (B) Overexpression of the dominant negative (DN) TOR (*TOR*^TED^) initially prevented destruction of the DEOM1s (arrow heads, +3h aHE), which occluded the DIOMs underneath (diagonal arrows) Subsequently (+12h, +20h), both DEOMs and DIOMs condensed and disintegrated. (C) Gene silencing of *Rm62* led to histolysis of DIOMs at +20h. (D) Silencing of *Cp1* induced degeneration of DIOMs in mid pupation from +38 to +70h. At +43h, the muscles in A2 completely histolysed (dashed arrows), while the DIOM1 (arrow) in the right hemi-segment of A3 disintegrated almost 20 hours later. Premature cell death of the three gene perturbations was restricted to larval persistent muscles. The DIOM1 in the left hemisegment of A3 survived (+75h, arrow). As can be seen at +75h, none of the gene perturbations caused discernible effects on the survival of newly formed adult muscle like dorsal abdominal muscles (double arrows) or the heart (heart shape). The scale bars in (D) represent 100 μm.

**Figure 5.**
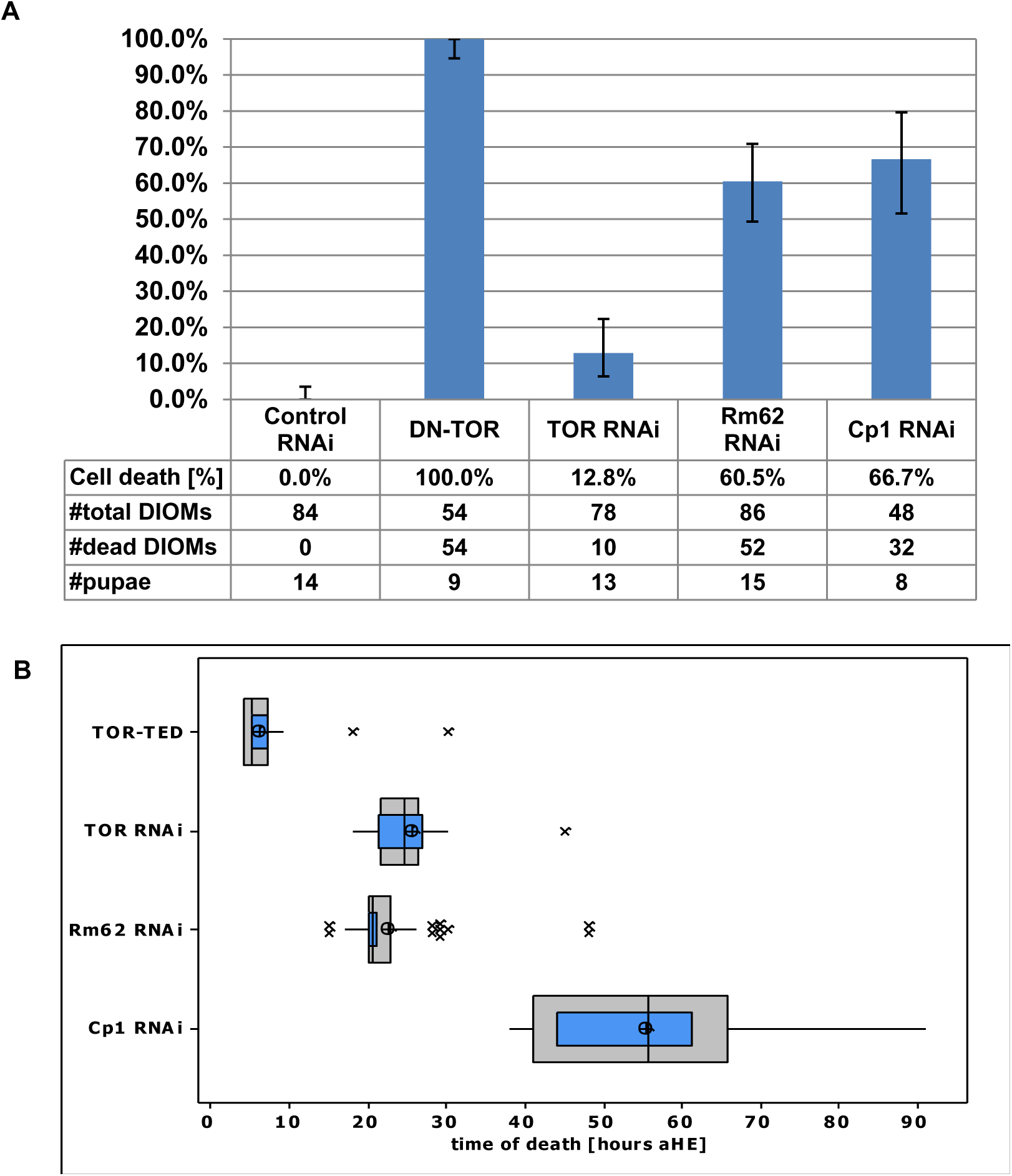
Selected gene perturbation cause cell death of DIOMs during different periods of pupation. (A) We scored proportion of DIOM1s (#cells) in abdominal segments A2 to A4 (6 per pupae) of the indicated number of samples (#pupae). Error bars indicate 95% confidence intervals of proportions. (B) Box-and-whisker plots show the distribution of time of death (TOD). Outer grey boxes show first quartiles, medians and third quartiles. Whiskers show minimum and maximum values. (*) symbols represent outliers. Inner blue boxes indicate 95% confidence intervals of the median. Circles indicate population means. The onset of muscle degeneration was assessed based on the earliest discernible alterations in cell morphology, e.g. loss of tubular shape. Cell death statistics were scored based on time-lapse images recorded by confocal microscopy at 22°C.

The silencing of the RNA helicase Rm62 induced premature histolysis during pupation in 60.5% of DIOM1s scored with a mean TOD of +22.4 h (Figure 4C, Figure 5). Muscle destruction gave rise to tau-GFP labelled sarcolytes, which were similar in size to those observed in control (Figure 4A) and considerably smaller than in *TOR*^TED^ overexpressing pupae (Figure 4B). As in control DEOM2s (Figure 6A), chromatin condensed prior to muscle fragmentation (Figure 6B). *Rm62* silencing did not delay or inhibit cell death of DEOM1s and DEOM2s. The results suggest that *TOR* and *Rm62* act cell-autonomously to protect persistent DIOMs from breakdown into sarcolytes.

**Figure 6.**
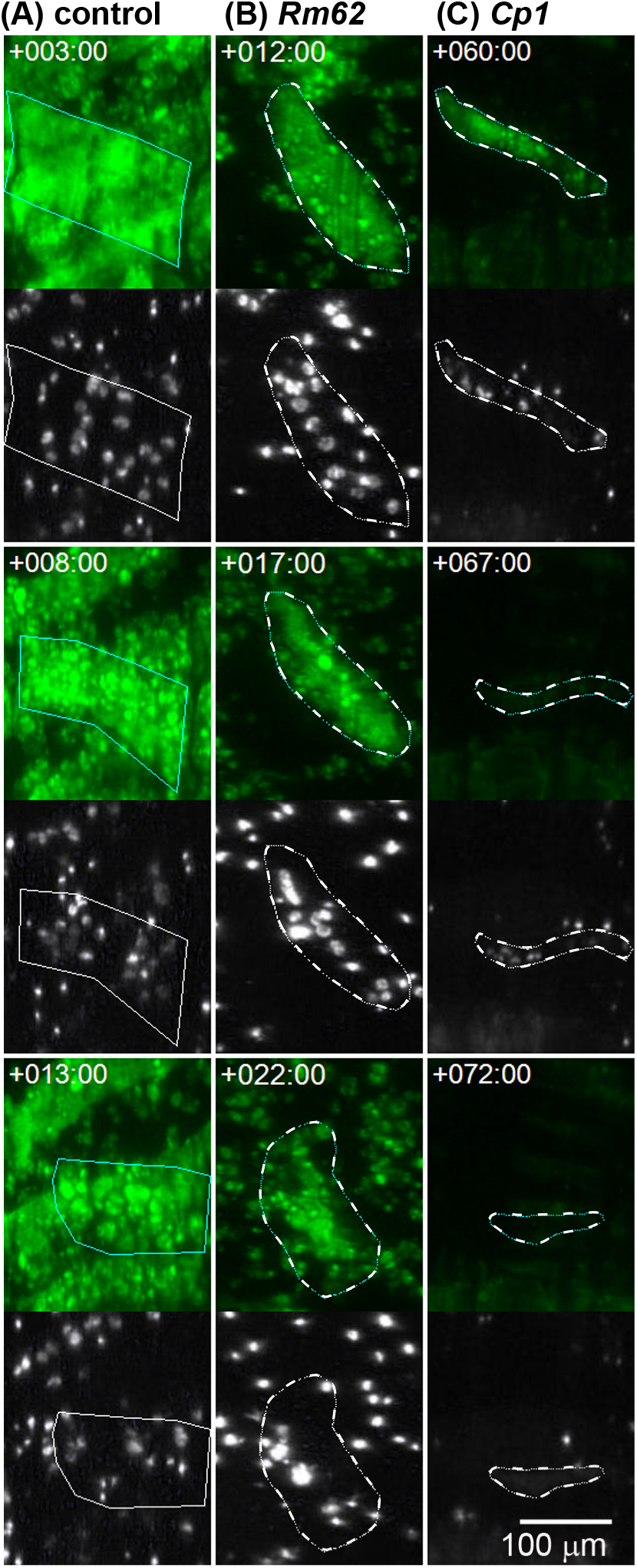
Different morphological changes during muscle cell death. (A-C) The top panels show muscle fibers labelled with tau-GFP (green) and outlined dotted lines. The bottom panels show the myonuclei labelled with histone-mKO (white). (A) A doomed DEOM2 expressing a control shRNA (see Figure 3A, ‘*’) disintegrates into fluorescently labelled vesicular fragments or sarcolytes. Nuclei become brighter as they condense. (B) A DIOM1 expressing *Rm62*-shRNA expressing (Figure 2C, arrow) undergoes premature histolysis similar to the PCD seen in the control DEOM2. (C) Cell death of *Cp1*-shRNA expressing muscles involves shrinking of cells and depletion of tau-GFP (compare +60 h and + 67h). Subsequently, discrete nuclear signals at +67h become fuzzy and weak at +72h.

### The Cathepsin-L· homolog Cp1 promotes DIOM survival in mid-pupation

Silencing of *Cp1*, the gene encoding the ortholog of the lysosomal Cathepsin-L·, led to decay of persistent muscles from +38 h onwards (mean TOD 55.3 ± 13.4 h) (Figure 5B), around one day later than in the case of *Rm62* RNAi (Figure 4D). In contrast to *TOR*^TED^ overexpression and *Rm62* RNAi, removal of DIOMs was not associated with the fragmentation of muscles into GFP-labelled sarcolytes. Instead, we observed cell shrinkage and a gradual decay of the cytoplasmic tau-GFP fluorescence, indicating proteolysis or leakage of the fluorescent protein (Figure 6C). Discrete nuclear histone-mKO labelling persisted slightly longer (3-6 hours) than tau-GFP, before eventually changing to hazy fluorescence of lower brightness, supporting the notion that *Cp1* RNAi, different from *TOR*^TED^ overexpression and *Rm62*-RNAi, induced a type of cell death that included rapid protein degradation. In conclusion, *Cp1* promotes survival of persistent muscles in mid-pupation. Compared to *TOR* and *Rm62, Cp1* suppresses a different type of cell death that involves cell shrinkage and rapid protein degradation instead of fragmentation.

### RNAi leads to loss of tubular morphology and degeneration of DIOMs

AMPK is a master regulator of energy metabolism that promotes energy (ATP) generating processes and inhibits energy consuming activities such as cell growth and proliferation [42]. In muscles, AMPK is activated during exercise by increasing AMP levels to promote ATP production and inhibit protein synthesis [43]. In the context of DIOM remodeling, AMPK could promote atrophy through the inhibition of protein synthesis. In early pupation between +13h to +20h aHE, when control muscle decreased in size (Figure 7A), *AMPKα* RNAi caused a loss of tubular morphology and decrease of tau-GFP fluorescence (Figure 7B; Additional file 4: Video 2). LSCM was performed at 25 °C. From +25h to +35h, DIOMs rounded up and displayed an increase in brightness of tau-GFP fluorescence. We will refer to the DIOM derived structures as muscle spheroids. The conversion of DIOMs to muscle spheroids showed 100% penetrance. The muscle spheroids floated within the pupal body, indicating a loss of attachment. Subsequently, muscle spheroids showed nuclear condensation (Figure 7C) and disappeared before eclosion. Due to their mobility, spheroids could not be tracked continuously in the field of view (Additional file 2: Video 2). Out of 47 spheroids scored in three pupae at +40h, we observed 7 nuclear condensation events between +35h to +44h. In late pupation at +70h, two remaining spheroid corresponded to an estimated ablation rate of 96%. As observed in 13 spheroids between +42h to +66h, the eliminations were accompanied by rapid depletions of tau-GFP within 30 minute intervals, while histone-mKO labeled condensed nuclei persisted for an additional 1-3 hours (Figure 7D). Similar to *Cp1* knockdowns, the destruction of *AMPKα* deficient muscle spheroids was not associated with the generation of sarcolytes. Despite the dramatic effects on DIOMs, other adult muscles did not display discernible defects and adult flies eclosed at wildtype rates.

**Figure 7.**
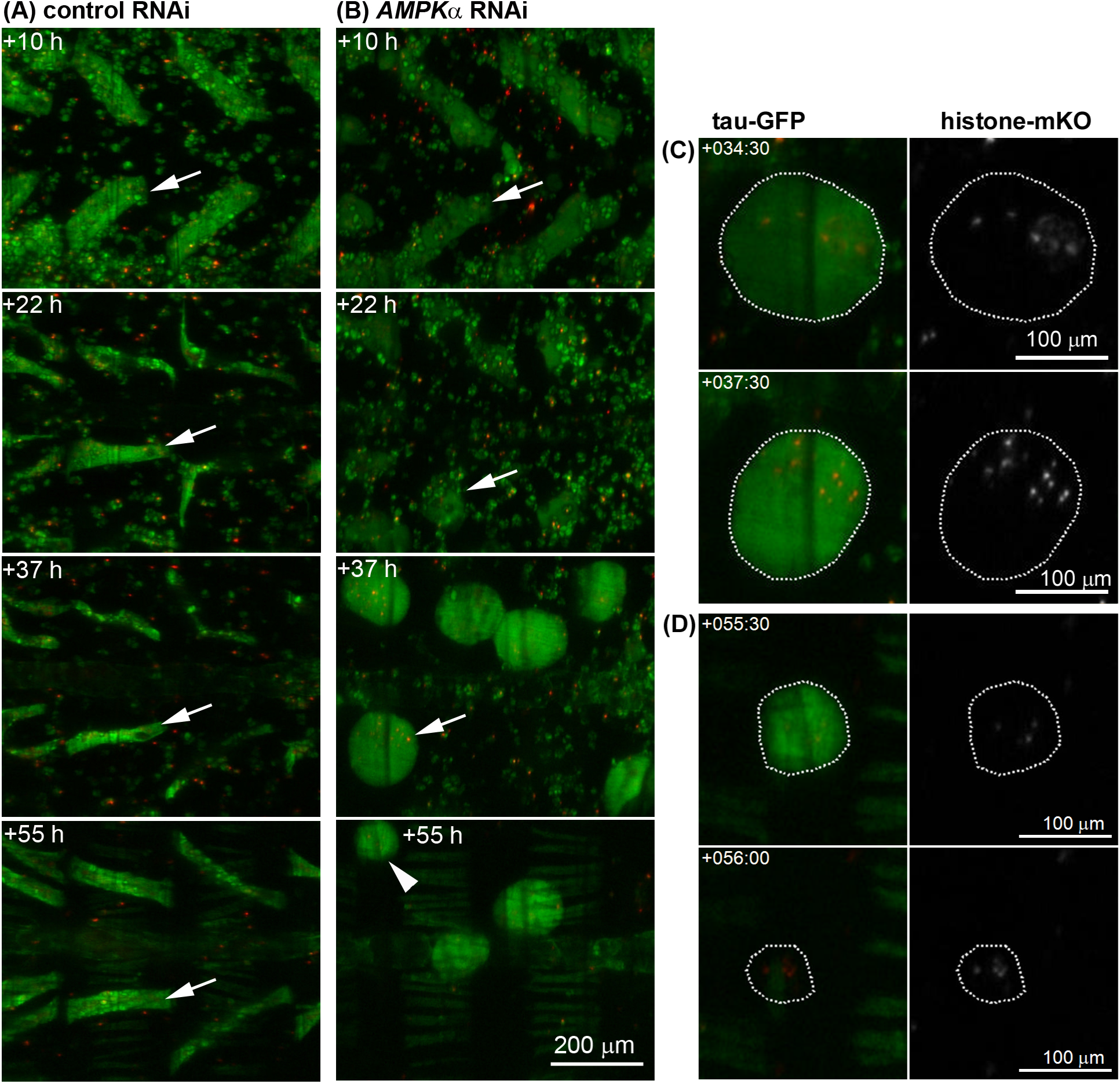
DIOMs detach and round up in response to *AMPKα* silencing. (A) Panels show the remodeling of DIOMs in abdominal segments A2 and A3. Remodeling involves reduction in diameter (atrophy) and rotation of the muscle fiber. (B) Silencing of *AMPKα* caused the loss of tubular morphology in DIOMs (arrow) between +10h to +22h. From +30 h onwards, DIOMs rounded up to give rise to floating muscle spheroids (+37h, see Video 3), indicating a loss of attachment. (C) *AMPKα*^shRNA^ muscle spheroids displayed condensation of histone-mKO tagged myonuclei (arrow in B, +37h). (D) In late pupation after +50h, muscle spheroids showed a sudden disappearance of tau-GFP, while condensed nuclei could still be observed for another 2 to 5 hours (arrow in B, +55h).

### Atrophin silencing inhibits histolysis of doomed muscles

Atrophin is a nuclear receptor co-repressor that has been shown to promote neurodegeneration [39]. *Atro* RNAi, similar to overexpression of DN-TOR and the N-terminal fragment of nuclear EAST [24], delayed the histolysis of DEOM1s (Figure 8). At +5h aHE, *Atro*^shRNA^ DEOM1s were intact, while their control counterparts were already shattered. In 7 *Atro*^shRNA^ pupae imaged by confocal microscopy, 79% of DEOM1s in A2-A4 (n=42 muscles, 95% CI: 63.2%, 89.7%) survived until +5h (Figure 9A). DEOM1 survival showed an anteriorposterior gradient, with 100%, 86% and 50% in A2, A3 and A4, respectively. Subsequent development of *Atro*^shRNA^ DEOM1s differed between abdominal segments (Figure 9B). While 80% of muscles (we scored 40 muscles per segment in 20 pupae using a macro zoom microscope) in A2 remained intact until the end of pupation, most DEOM1s in the 3^rd^ and all in the 4^th^ and 5^th^ segment shrank and broke apart into sarcolytes in the next 5 to 7 hours (Figure 8B; Additional file: Video 3). Therefore, *Atro* RNAi is the first known gene perturbation that is able to transform a doomed to a persistent muscle. We also observed rescue of DEOM1s in the first abdominal segment, which we were unable to accurately quantify due to limited visibility of muscles. DEOM1s protected from histolysis did not show nuclear condensation (Figure 8D). Histolysis of DEOM2s did not show differences between control and *Atro* RNAi, indicating that silencing of *Atro* alone is not sufficient to delay or block the cell death of doomed muscles. *Atro* RNAi induced the transient formation of large vacuoles that were not seen in controls (Figure 8B; +12h). Furthermore, *Atro* silencing also caused a segment-specific loss of DIOM1s (Figure 9C), which contrary to the histolysis in inhibition in DEOM1s rescue showed a posterior to anterior gradient. While 53% of DIOM1s vanished in A5 and 15% in A4, no cell death was observed in segments A2 and A3.

**Figure 8.**
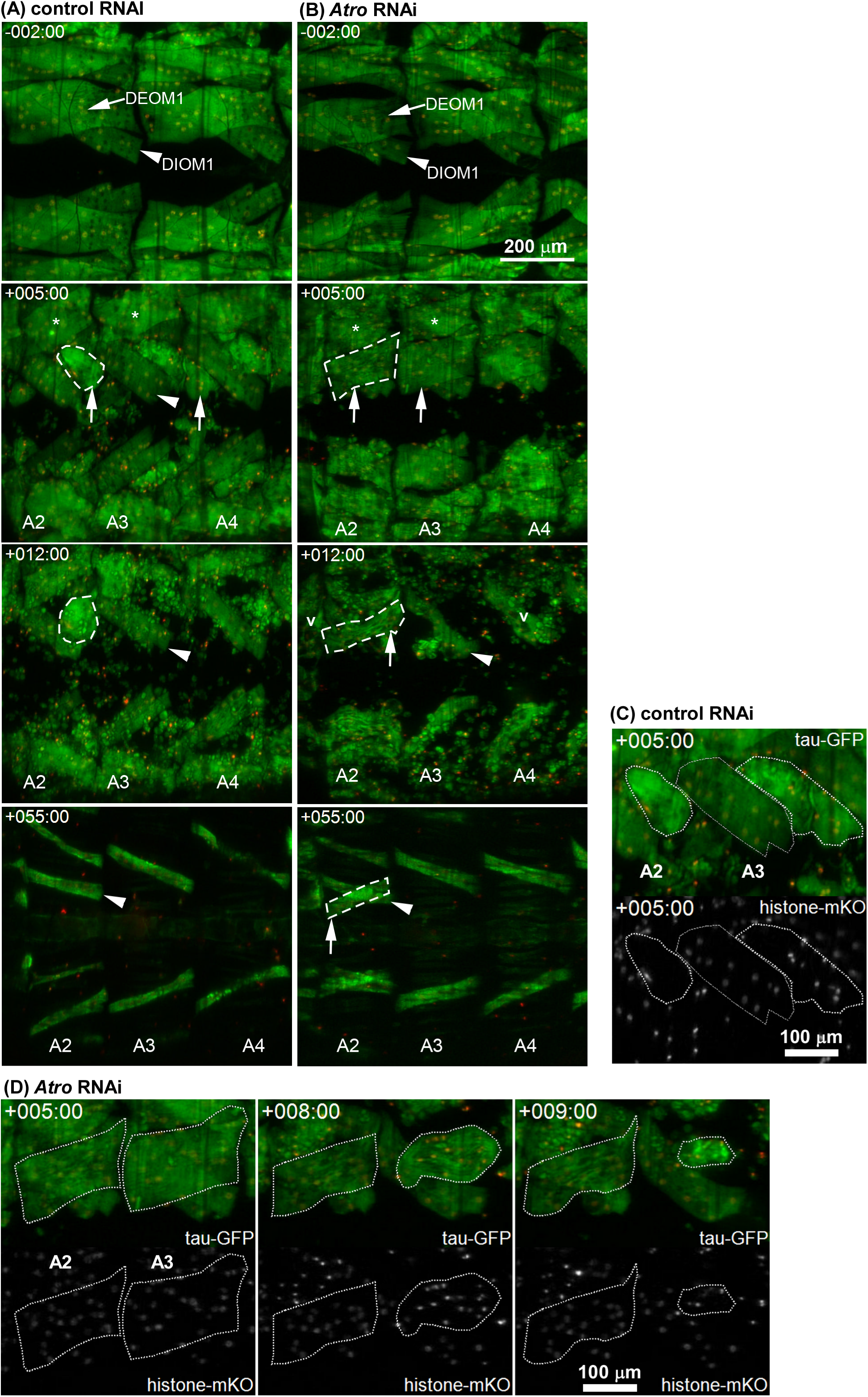
Silencing of *Atrophin* inhibits muscle histolysis. (A) In controls during HE, DEOM1s (arrow,-2h) disintegrated into sarcolytes (+5h, arrows), revealing the DIOM1s (arrow head, +5h) underneath. The more laterally located DEOM2s (*, +5h) histolyzed later after HE. (B) Histolysis of *Atro*^shRNA^ DEOM1s was delayed until early pupation (arrows, +5h), resulting in intact muscles (arrows) at +5h. Subsequently, DEOM1s in A3-A5 were destroyed. The DEOM1s in A2 persisted into adulthood, resulting in crosswise overlapping muscles at +55h. *Atro* silencing led to transient formation of vacuoles (v) at +12h. (C) Histolysis of control DEOM1s (white contours, compare to A, +5h) was accompanied by chromatin (histone-mKO) condensation. Compare to myonuclei in DIOM1 (grey contour). (D) Delayed DEOM1 histolysis in A3 was accompanied by cell shrinkage and chromatin condensation, which did not occur in the corresponding muscle in A2. Time-lapse datasets were recorded at 25°C.

**Figure 9.**
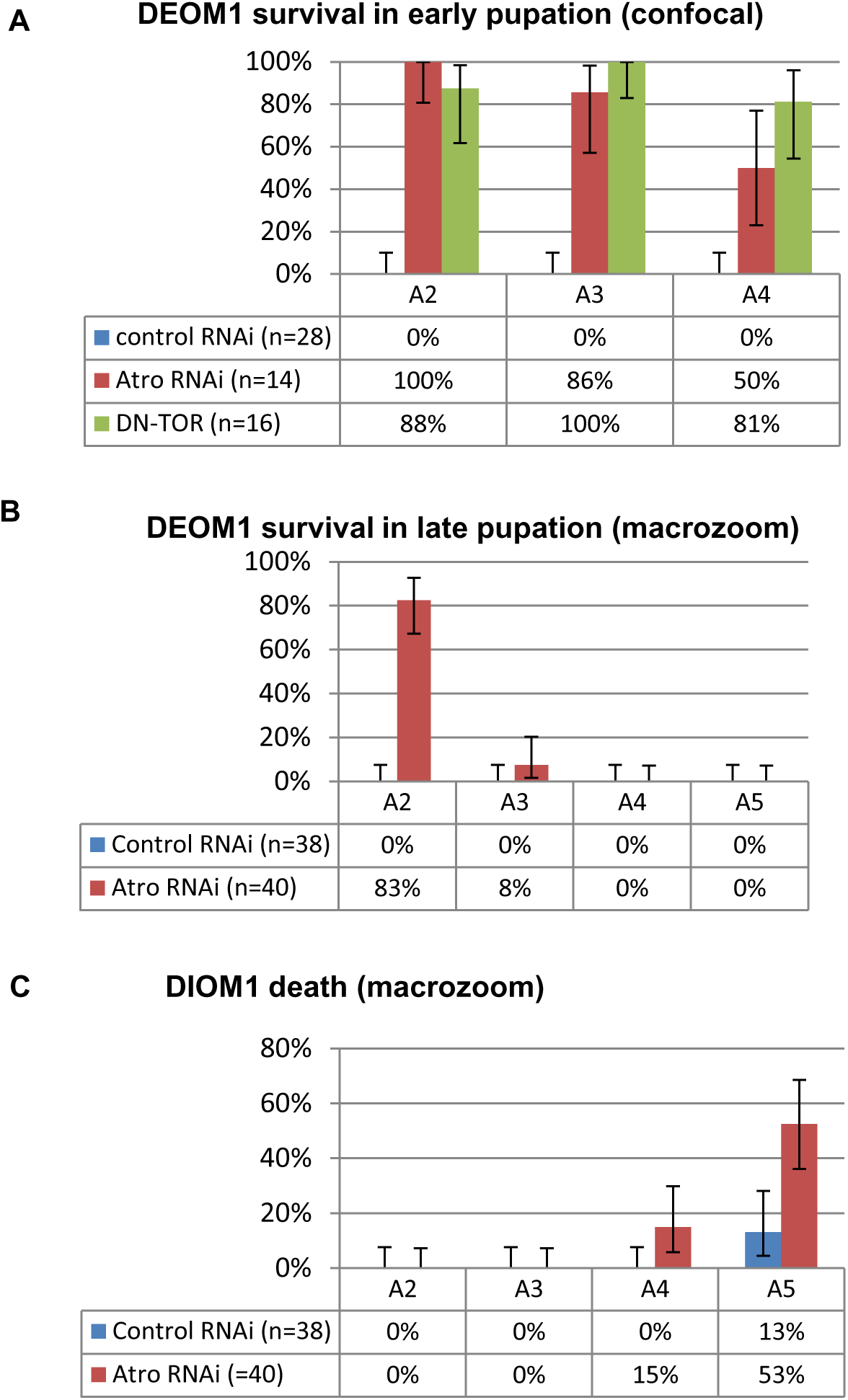
*Atrophin* RNAi inhibits histolysis of DEOMs. **(A)** Survival rates of doomed (DEOM1s) and (B) cell death rates of persistent muscles (DIOM1s) were scored for abdominal segments A2-A5 in late pupation. We examined 19 control and 20 *Atro* shRNA expressing pupae by macro zoom microscopy. As such, we scored 38 control and 40 *Atro* RNAi muscles per segment. Errors bars indicate 95% confidence intervals of the proportions.

## Discussion

Although larval body wall muscles are structurally and physiologically similar, they respond differently to changing environmental conditions during metamorphosis. Our method provided new insights into the genes that determine if, when and how muscles execute programmed self-destruction (Figure 10). Due to its ability to characterize dynamic and transient phenotypes, time-lapse image analysis allowed us to differentiate gene perturbations based on four main criteria; (1) whether they promoted or inhibited cell death, (2) which muscles were affected, (3) the developmental period of phenotypic abnormalities and (4) the morphological transformations during muscle destruction.

**Figure 10.**
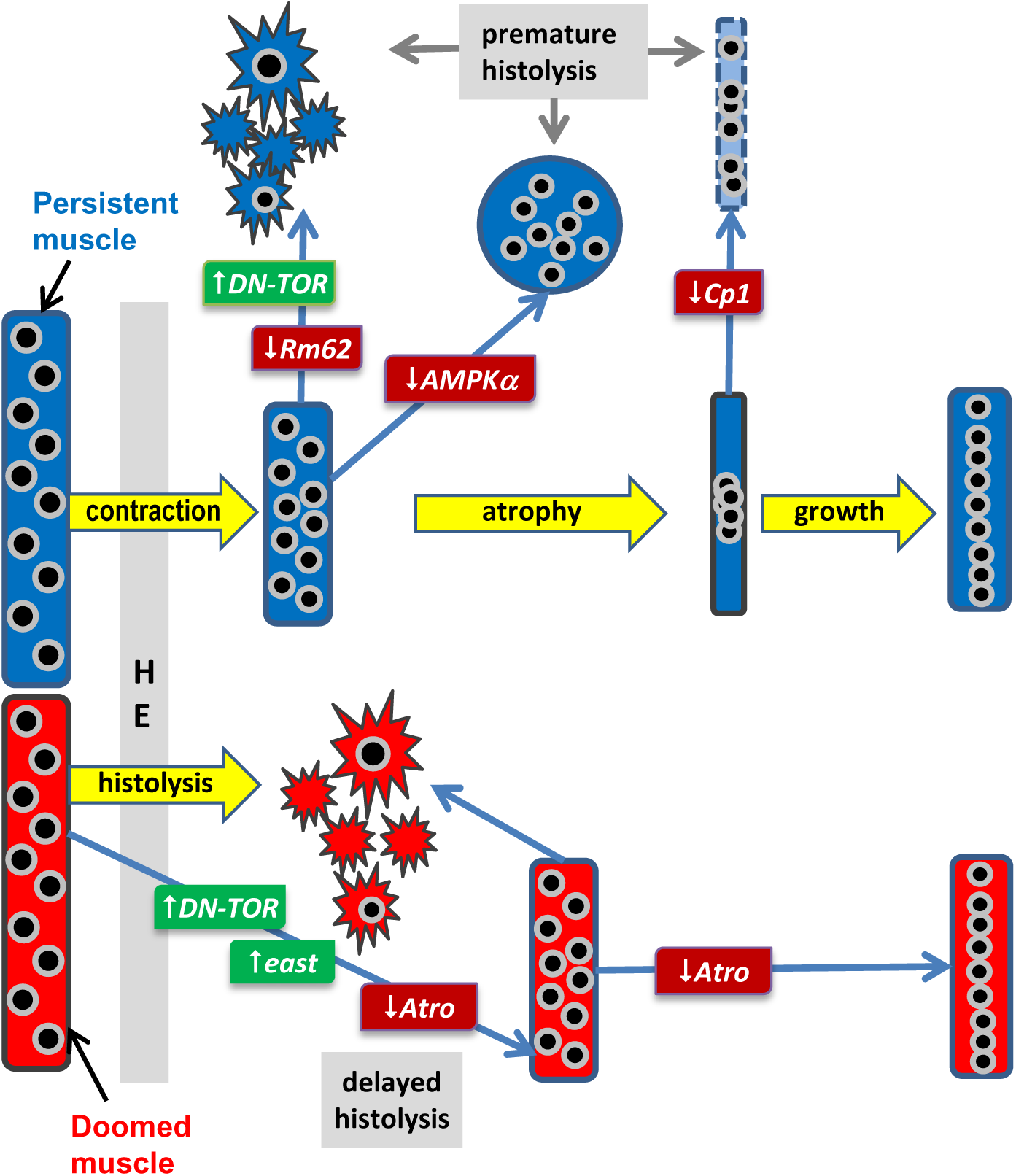
Gene perturbations that affect cell death and survival of larval abdominal muscles during metamorphosis. Larval persistent muscles undergo remodeling to adult temporary muscles, which involves atrophy, nuclear migration and hypertrophy. We identified gene perturbations that specifically cause premature cell death of persistent muscles without affecting the survival of newly formed adult muscles. *TOR* and *Rm62* protect DIOMs from fragmentation into sarcolytes in early pupation. In mid-pupation, *Cp1* prevents a different type of cell death that involves leakage and/or lysis of fluorescent proteins. Silencing of *AMPKα* induces detachment, loss of tubular morphology and ablation of muscle spheroids. Doomed muscles undergo histolysis during and right after HE, which can be delayed by overexpression of dominant negative (DN) TOR, a fragment of the *east* gene and *Atro* RNAi. In addition, *Atro* RNAi is the only gene perturbation so far that promotes survival of a subset of DEOMs until adulthood.

In early pupation, *TOR*^TED^ *overexpression* and *Rm62*-RNAi induced the breakdown of DIOMs into sarcolytes. The phenotypic similarity to normal histolysis of DEOMs suggests that *TOR* and *Rm62* cell autonomously inhibit cell death of persistent muscles. Evidence in the literature indicates that mTOR can either promote or inhibit apoptosis depending on the biological context [39], supporting the idea that *TOR* may protect remodeled muscles from histolysis. However, since *TOR*^TED^ overexpression may also create unphysiological conditions, we cannot rule out the possibility that the phenotype is an artefact. Circumstantial evidence links the RNA helicase Rm62 to TOR signaling since Rm62 protein was detected in the same complex as the protein Poly, which is a *Drosophila* homolog of a transcription elongation factor Elp6 and acts as a positive regulator of InR-TOR signaling [40]. Time-lapse analysis uncovered phenotypic differences between the two gene perturbations. Compared to *TOR*^TED^, *Rm62*-RNAi induced histolysis occurred 15 hours later and did not delay histolysis of DEOMs. In the context of abdominal muscles, *TOR*^TED^ did not exactly behave like a dominant negative version of the endogenous TOR protein since TOR-RNAi mainly affected cell size and had negligible effects on cell survival [30]. We can only speculate that the regulation of cell survival and cell growth are mediated by different protein complexes.

Cp1, the fly ortholog of lysosomal Cathepsin-L, has been proposed to mediate autophagic cell death since its gene expression is upregulated during salivary gland cell death [44]. Contrary to this prediction, our results showed that Cp1 acts as a suppressor of cell death in DIOMs. In contrast to the phenotypes caused by *TOR*^TED^ overexpression and *Rm62* RNAi, cell death occurred in a later stage of metamorphosis and was not associated with the formation of sarcolytes. Prior to DIOM disappearance, fluorescence of the tau-GFP and histone-mKO reporter decreased and became diffuse, suggesting that *Cp1*-RNAi induced protein and chromatin degradation inside muscles rather than their fragmentation into sarcolytes. A plausible explanation for the loss-of-function phenotype is that Cp1 may act to break down PCD-promoting proteins that start to accumulate in persistent muscles towards late metamorphosis, thus ensuring that DIOMs only degenerate after eclosion.

Previous studies have shown that autophagy promoted cell death in tissues like the midgut in *Drosophila* metamorphosis [22]. Since *TOR*^TED^ is a potent inducer of autophagy in the fat body [41], activation of autophagy may trigger cell death of wildtype DEOMs and *TOR*^TED^ overexpressing DIOMs. In a parallel study, we found that RNAi of 5 autophagy related genes (*Atg5*, *Atg9*, *Atg12*, *Atg17*, and *Atg18*) caused inhibition of atrophy during the remodeling of DIOMs [30]. We examined 25 time-lapse datasets of pupae expressing shRNAs against Atg9 (7) and Atg5, Atg12 and Atg18 (6 each). However, we unable to find evidence that silencing of these genes inhibited or blocked histolysis of DEOMs, as we demonstrated for Atro-RNAi and *TOR*^TED^ overexpression. Hence, in agreement with a previous study [23], autophagy does not appear to play a role in mediating cell death of larval muscles.

Previous studies have shown that ecdysone receptor signaling cell-autonomously triggers apoptosis in DEOMs [23]. Our results corroborate previous reports in rats and moths, which concluded that PCD in muscles did not display a variety of morphological features observed in classical apoptosis, such as phagocytosis or DNA fragmentation [10, 17]. Although we observed chromatin condensation prior to and during histolysis, we did not find evidence for nuclear fragmentation that can be observed in mono-nuclear non muscle cells. In addition, the intensity and localization pattern of histone-mKO in sarcolytes showed remarkable stability, arguing against strong proteolytic activities which are commonly associated with apoptosis. Similarly, non-nuclear fusion proteins like tau-GFP, Cp1-mKO2 and Pros35-mOrange2 provided stable fluorescent labeling of sarcolytes. Our results are not necessarily in conflict with studies in *Manduca* which showed that PCD in ISMs involved the increased activation of the autophagy-lysosomal and ubiquitin-proteasome pathways [14]. A tight control over proteolysis and, probably, lipolysis may ensure that sarcolytes continue to function as amino acid reservoirs that help to feed the developing fly during pupation.

So far, little is known about the genes that mediate DEOM PCD downstream of the ecdysone receptor. Our results suggest that *Atrophin* may play a role in promoting muscle degeneration. Consistent with previous reports that *Atro* overexpression leads to neuronal cell death [39, 45], our data show that *Atro* knockdown can inhibit histolysis of doomed larval muscles. In both neurons and muscles, *Atro* appears to show genetic interactions with TOR signaling. While *TOR*^TED^ overexpression enhances neurodegeneration elicited by *Atro* gain-of-function [45], both Atro-RNAi and *TOR*^TED^ overexpression delay histolysis of DEOM1s.

The response of DEOMs to *Atro* silencing was cell-specific. Histolysis of all DEOM1s was delayed, while destruction of DEOM2s occurred at the normal time. Moreover, a subset of DEOM1s, particularly in the anterior abdominal segments was resistant to histolysis and survived until the end of metamorphosis. The reason for this specificity remains to be elucidated. Since Atro protein acts as a transcriptional co-repressor that recruits histone deacetylases [46], it may repress the transcription of pro-survival genes in doomed muscles.Given that loss of *Atro* is not sufficient to block PCD in all DEOMs, other redundant transcriptional regulators may repress yet to be discovered target genes.

## Conclusions

Live microscopy of *Drosophila* metamorphosis is a powerful tool to study the processes and genes that promote and prevent cell death in muscles. Candidates of protective genes include *Rm62, Cp1, TOR* and *AMPKα*, all of which were discovered in a pilot screen involving 98 preselected unique genes. Time-lapse image analysis provides detailed insights into muscle degeneration that would be difficult to obtain from the study of fixed tissues. Cellular parameters like chromatin condensation, size, shape and fragmentation can be continuously monitored. For different genotypes, we can compare the time of death of different populations of muscles. Using a pilot screen, we identified 5 gene perturbations that could delay or block cell death of doomed muscles, or induce premature histolysis of persistent muscles. *Atrophin* is so far the only gene, whose silencing can rescue a subset of DEOMs until the end of metamorphosis. Remodeled muscles undergo atrophy. Interestingly, none of the perturbations that induced cell death of remodeled muscles showed discernible effects on the survival of newly formed adult muscles, suggesting that this approach may help find genes that may render muscles undergoing atrophy more susceptible to degeneration. Unbiased genome-wide screen in the future promise to identify more genes involved in myogenic PCD during insect metamorphosis, some of which may be evolutionarily conserved and relevant for human muscle wasting.

## List of abbreviations

aHE: after head eversion;
AMPKα: AMP-activated protein kinase α subunit;
Atro: Atrophin;
Cp1: Cysteine proteinase 1;
DEOM: dorsal external oblique muscle;
DIOM: dorsal internal oblique muscles;
DN: dominant negative;
GFP: green fluorescent protein;
HE: head eversion;
ISM: intersegmental muscle;
LSCM: laser scanning confocal microscopy;
MIP: maximum intensity projection;
mKO: monomeric kusabira orange;
mTOR: mammalian target of rapamycin;
PCD: programmed cell death;
PPT: prepupal to pupal transition;
RNAi: RNA interference;
sh: small hairpin;
TOD: time of death;
TOR^TED^: target of rapamycin toxic effector domain.

## Authors’ contributions

M.W. conceived the study and designed the experiments. Y.K., W.C.P. and M.W. performed the experiments and analysed the data. M.W. drafted the article.

## Acknowledgements

M.W. and W.C.P. were affiliated with the Bioinformatics Institute (BII) until 2013, YK until 2015. We thank the TRiP at Harvard Medical School (NIH/NIGMS R01-GM084947) for providing transgenic RNAi fly stocks used in this study. We thank the Bloomington Drosophila stock centre for providing fly stocks. We thank Hwee Kuan Lee and Lin Feng for providing academic advice to YK. We thank the IMCB (ACC and SC labs) for the use of Drosophila facilities.

## Additional files

**Additional file 1: Table S1.** List of gene perturbations tested for muscle defects in pilot screen.

**Figure S1.**
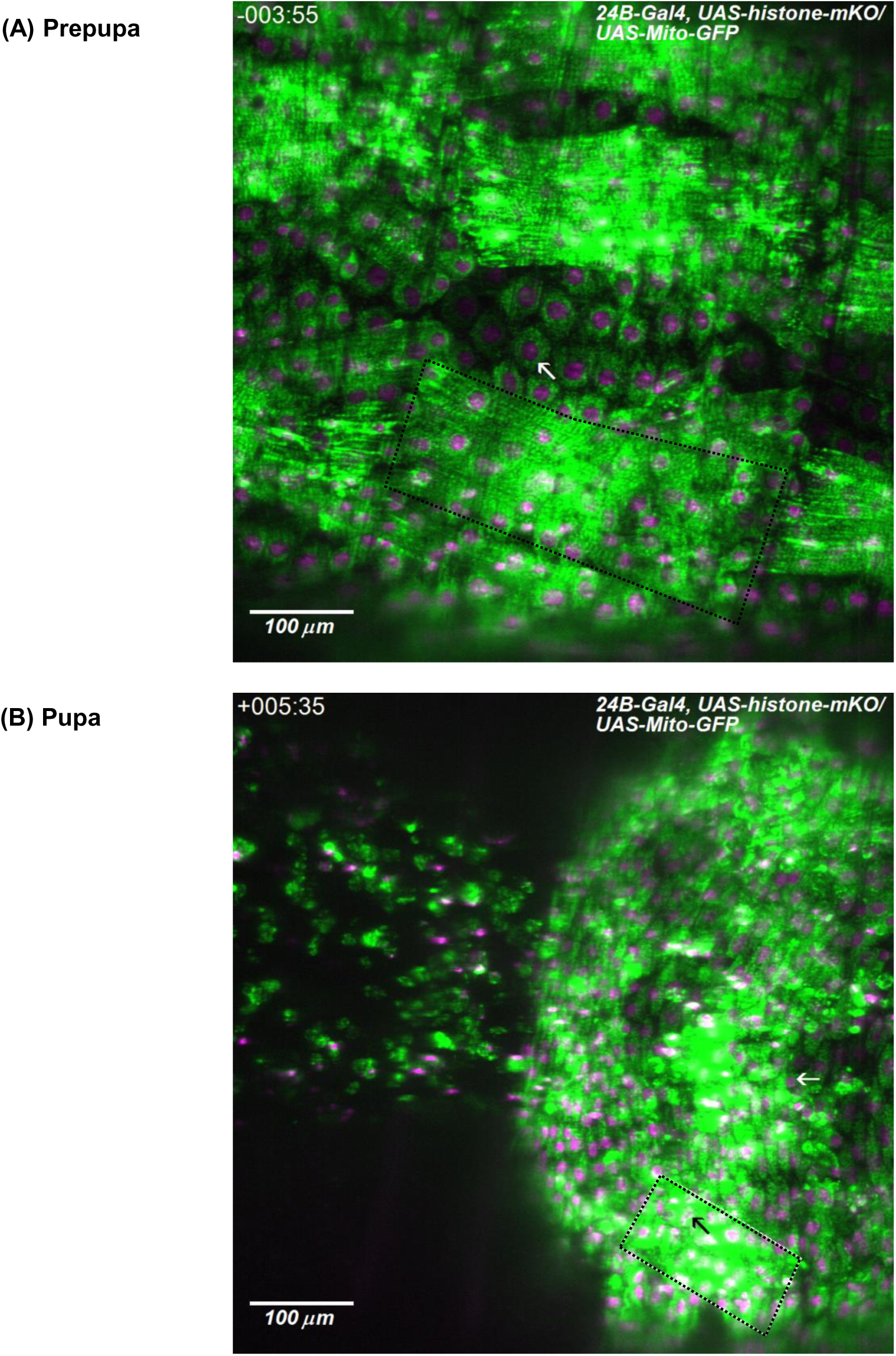
Additional file 2: Figure S1. Expression of the 24B-GAL4 driver in metamorphosis. In (A) prepupae and (B) pupae, the mesodermal 24B-Gal4 driver is expressed in multinucleated muscles (outlined in black) and more apically located mono-nucleated cells (white arrow). Cells were labelled with the mitochondrial marker UAS-Mito-GFP (green) and UAS-histone-mKO (magenta). (B) During pupation, mono-nucleated cells (white arrow) undergo PCD and show nuclear fragmentation, while nuclei (black arrow) in muscles undergoing histolysis, like the DIOM2s (outlined in black), condense without showing fragmentation (see Figure 2).

**Figure S2.**
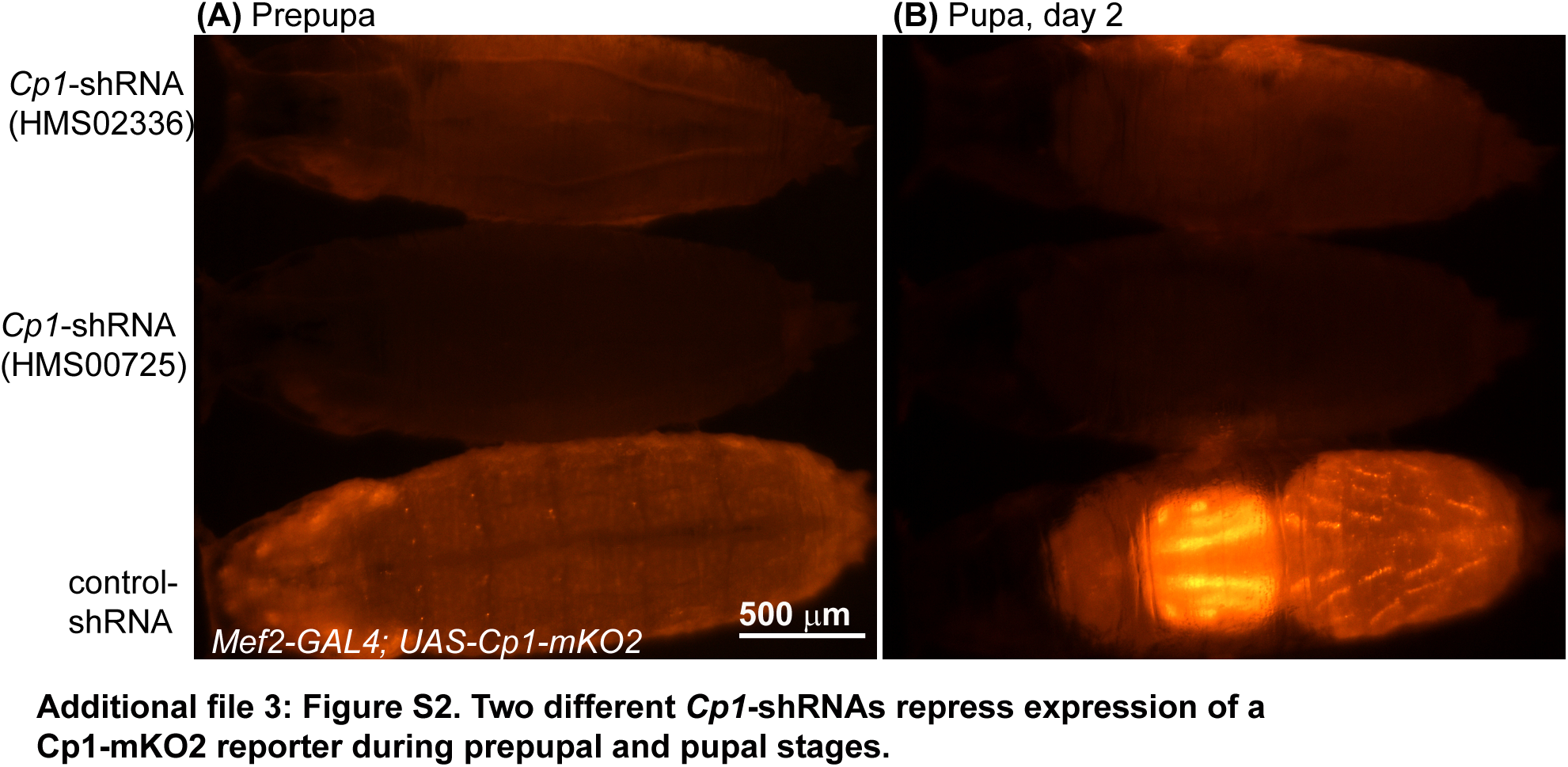
Additional file 3: Figure S2. Two different Cp1-shRNAs silence expression of a Cp1-mKO2 reporter during prepupal (A) and pupal (B) stages. The construct HMS00725 (second row) was used in most experiments.

**Additional file 4: Video 1. Premature cell death of persistent muscles. ([YouTube](https://youtu.be/k6lPTPLl9ks))**Overexpression of dominant negative TOR (*TOR*^TED^), and RNAi of *Rm62* and *Cp1* induce histolysis of DIOMs, which persist throughout metamorphosis until adulthood in controls (top left). The time-lapse frames show the development from the prepupal (-6 h) to the late pupal stage (+100 h). Muscles are labelled with tau-GFP (green) and histone-mKO (red). Pupae are oriented in an anterior-posterior direction from left to right. Views show part of the thorax and abdominal segments A1 to A4. Note the stability of fluorescently labelled sarcolytes derived from histolyzed muscles. None of the gene perturbations caused obvious cell death of newly formed adult muscles, such as IFMs, heart and abdominal muscles. The time-lapse images were recorded at 22 °C.

**Additional file 5: Video 2. Silencing of *AMPKα* leads to loss of tubular morphology and disintegration of DIOMs. ([YouTube](https://youtu.be/JLHbiFsojPA))**Compared to control muscles (left panel),*AMPKα*^*shRNA*^ DIOMs gradually lose tubular morphology from +15h aHE onwards. From around +30h, DIOMs turn into floating muscle spheroids, indicating a loss of cell attachment. Later, green fluorescence of spheroids rapidly disappears within 30 minute intervals (see arrow at +56:30h), while condensed nuclei stay behind, indicating rapid lysis without formation of sarcolytes. The time-lapse images were recorded at 25 °C. Muscles were labeled with Mhc-tau-GFP (green) and histone-mKO (magenta).

**Additional file 6: Video 3. Silencing of *Atrophin* inhibits muscle histolysis in metamorphosis. ([YouTube](https://youtu.be/aG9JB8HZNdc))**Atro-RNAi (right panel) delays histolysis of DEOM1s in all abdominal segments. While control DEOMs (left panels) are destroyed at the onset of HE, *Atro*^shRNA^ muscles persist for up to 10 hours into pupation. In abdominal segment A2,DEOMs (arrow) survive until adulthood. The time-lapse images were recorded at 25 °C. Muscles were labeled with Mhc-tau-GFP (green) and histone-mKO (magenta). The scale bars correspond to 200 μm.

## References

1. Evans WJ: Skeletal muscle loss: cachexia, sarcopenia, and inactivity. Am J Clin Nutr 2010, 91:1123S–1127S.

2. Nowak KJ, Davies KE: Duchenne muscular dystrophy and dystrophin: pathogenesis and opportunities for treatment. EMBO Rep 2004, 5:872–876.

3. Jungbluth H, Gautel M: Pathogenic mechanisms in centronuclear myopathies. Front Aging Neurosci 2014, 6:339.

4. Laplante M, Sabatini DM: mTOR signaling in growth control and disease. Cell 2012, 149:774–793.

5. Elkina Y, von Haehling S, Anker SD, Springer J: The role of myostatin in muscle wasting: an overview. J Cachexia Sarcopenia Muscle 2011, 7.143–151

6. Sandri M: Autophagy in skeletal muscle. FEBS Lett 2010, 584:1411–1416.

7. Polge C, Attaix D, Taillandier D: Role of E2-Ub-conjugating enzymes during skeletal muscle atrophy. Front Physiol 2015, 6:59.

8. Fulle S, Sancilio S, Mancinelli R, Gatta V, Di Pietro R: Dual role of the caspase enzymes in satellite cells from aged and young subjects. Cell Death Dis 2013, 4:e955.

9. Xiao R, Ferry AL, Dupont-Versteegden EE: Cell death-resistance of differentiated myotubes is associated with enhanced anti-apoptotic mechanisms compared to myoblasts. Apoptosis 2011, 16: 221–234.

10. Borisov AB, Carlson BM: Cell death in denervated skeletal muscle is distinct from classical apoptosis. Anat Rec 2000, 258:305–318.

11. Bruusgaard JC, Egner IM, Larsen TK, Dupre-Aucouturier S, Desplanches D, Gundersen K: No change in myonuclear number during muscle unloading and reloading. J Appl Physiol 2012, 113:290–296.

12. Nishikawa A, Hayashi H: Spatial, temporal and hormonal regulation of programmed muscle cell death during metamorphosis of the frog Xenopus laevis. Differ Res Biol Divers 1995, 59:207–214.

13. Finlayson LH: Normal and Induced Degeneration of Abdominal Muscles during Metamorphosis in the Lepidoptera. Q j Microsc Sci 1956, s3-97:215–233.

14. Schwartz LM: Atrophy and programmed cell death of skeletal muscle. Cell Death Differ 2008, 15:1163–1169.

15. Lockshin RA, Williams CM: Programmed cell death—II. Endocrine potentiation of the breakdown of the intersegmental muscles of silkmoths. j Insect Physiol 1964, 10:643–649.

16. Jones ME, Schwartz LM: Not all muscles meet the same fate when they die. Cell Biol Int 2001, 25:539–545.

17. Schwartz LM, Smith SW, Jones ME, Osborne BA: Do all programmed cell deaths occur via apoptosis? Proc Natl Acad Sci U S A 1993, 90:980–984.

18. Haralalka S, Abmayr SM: MyoNast fusion in Drosophila. Exp Cell Res 2010, 316:3007–3013.

19. Demontis F, Perrimon N: integration of Insulin receptor/Foxo signaling and dMyc activity during muscle growth regulates body size in Drosophila. Development 2009, 136:983–993.

20. Wasser M, Bte Osman Z, Chia W: EAST and Chromator control the destruction and remodeling of muscles during Drosophila metamorphosis. Dev Biol 2007, 307.380–393,

21. Kimura KI, Truman JW: Postmetamorphic cell death in the nervous and muscular systems of Drosophila melanogaster. J Neurosci 1990, 10:403–401.

22. Denton D, Shravage B, Simin R, Mills K, Berry DL, Baehrecke EH, Kumar S: Autophagy, not apoptosis, is essential for midgut cell death in Drosophila. Curr Biol 2009, 19:1741–1746.

23. Zirin J, Cheng D, Dhanyasi N, Cho J, Dura J-M, Vijayraghavan K, Perrimon N: Ecdysone signaling at metamorphosis triggers apoptosis of Drosophila abdominal muscles. Dev Biol 2013, 383:275–284.

24. Chinta R, Tan JH, Wasser M: The study of muscle remodeling in Drosophila metamorphosis using in vivo microscopy and bioimage informatics. BMC Bioinformatics 2012, 13 Suppl 17:S14.

25. Brand AH, Perrimon N: Targeted gene expression as a means of altering cell fates and generating dominant phenotypes. Development 1993, 118:401–415.

26. Schnorrer F, Schönbauer C, Langer CCH, Dietzl G, Novatchkova M, Schernhuber K, Fellner M, Azaryan A, Radolf M, Stark A, Keleman K, Dickson BJ: Systematic genetic analysis of muscle morphogenesis and function in Drosophila. Nature 2010, 464:287–291.

27. Bour BA, O'Brien MA, Lockwood WL, Goldstein ES, Bodmer R, Taghert PH, Abmayr SM, Nguyen HT: Drosophila MEF2, a transcription factor that is essential for myogenesis. Genes Dev 1995, 9:730–741.

28. Chen EH, Olson EN: Antisocial, an intracellular adaptor protein, is required for myoblast fusion in Drosophila. Dev Cell 2001, 1:705–715.

29. Ni J-Q, Zhou R, Czech B, Liu L-P, Holderbaum L, Yang-Zhou D, Shim H-S, Tao R, Handler D, Karpowicz P, Binari R, Booker M, Brennecke J, Perkins LA, Hannon GJ, Perrimon N: A genome-scale shRNA resource for transgenic RNAi in Drosophila. Nat Methods 2011, 8:405–407.

30. Kuleesha J, Puah WC, Wasser M: A model of muscle atrophy based on live microscopy of muscle remodelling in Drosophila metamorphosis. R Soc Open Sci 2016, 3:150–517.

31. Puah WC, Wasser M: Live imaging of muscles in Drosophila metamorphosis: Towards highthroughput gene identification and function analysis. Methods 2015.

32. Pilling AD, Horiuchi D, Lively CM, Saxton WM: Kinesin-1 and Dynein Are the Primary Motors for Fast Transport of Mitochondria in Drosophila Motor Axons. Mol Biol Cell 2006, 17:2057–2068.

33. Kuleesha Y, Puah WC, Lin F, Wasser M: FMAJ: a tool for high content analysis of muscle dynamics in Drosophila metamorphosis. BMC Bioinformatics 2014, 15 Suppl 16:S6.

34. Puah WC, Cheok LP, Biro M, Ng WT, Wasser M: TLM-Converter: reorganization of long timelapse microscopy datasets for downstream image analysis. BioTechniques 2011, 51:49-50, 52–53.

35. Freelmage [http://freeimage.sourceforge.net/index.html]

36. libics [http://libics.sourceforge.net/]

37. Zeng X, Han L, Singh SR, Liu H, Neumüller RA, Yan D, Hu Y, Liu Y, Liu W, Lin X, Hou SX: Genomewide RNAI screen identifies networks involved in intestinal stem cell regulation in Drosophila. Cell Rep 2015, 10:1226–1238.

38. Sopko R, Foos M, Vinayagam A, Zhai B, Binari R, Hu Y, Randklev S, Perkins LA, Gygi SP, Perrimon N: Combining genetic perturbations and proteomics to examine kinase-phosphatase networks in Drosophila embryos. Dev Cell 2014, 31:114–127.

39. Karres JS, Hilgers V, Carrera I, Treisman J, Cohen SM: The conserved microRNA miR-8 tunes atrophin levels to prevent neurodegeneration in Drosophila. Cell 2007, 131:136–145.

40. Dobi KC, Halfon MS, Baylies MK: Whole-Genome Analysis of Muscle Founder Cells Implicates the Chromatin Regulator Sin3A in Muscle Identity. Cell Rep 2014, 8:858–870.

41. Scott RC, Schuldiner O, Neufeld TP: Role and regulation of starvation-induced autophagy in the Drosophila fat body. Dev Cell 2004, 7:167–178.

42. Hardie DG, Ross FA, Hawley SA: AMPK: a nutrient and energy sensor that maintains energy homeostasis. Nat Rev Mol Cell Biol 2012, 13:251–262.

43. Lantier L, Fentz J, Mounier R, Leclerc J, Treebak JT, PehmϘller C, Sanz N, Sakakibara I, Saint-Amand E, Rimbaud S, Maire P, Marette A, Ventura-Clapier R, Ferry A, Wojtaszewski JFP, Foretz M, Viollet B: AMPK controls exercise endurance, mitochondrial oxidative capacity, and skeletal muscle integrity. FASEB J Off Publ Fed Am Soc Exp Biol 2014, 28:3211–3224.

44. Gorski SM, Chittaranjan S, Pleasance ED, Freeman JD, Anderson CL, Varhol RJ, Coughlin SM, Zuyderduyn SD, Jones SJM, Marra MA: A SAGE approach to discovery of genes involved in autophagic cell death. Curr Biol 2003, 13:358–363.

45. Nisoli I, Chauvin JP, Napoletano F, Calamita P, Zanin V, Fanto M, Charroux B: Neurodegeneration by polyglutamine Atrophin is not rescued by induction of autophagy. Cell Death Differ 2010, 17: 1577–1587.

46. Zhang Z, Feng J, Pan C, Lv X, Wu W, Zhou Z, Liu F, Zhang L, Zhao Y: Atrophin-Rpd3 complex represses Hedgehog signaling by acting as a corepressor of CiR. J Cell Biol 2013, 203:575–583.

